# A synthetic delivery vector for mucosal vaccination

**DOI:** 10.1101/2023.05.15.540756

**Authors:** Anne Billet, Justine Hadjerci, Thi Tran, Pascal Kessler, Jonathan Ulmer, Gilles Mourier, Marine Ghazarian, Anthony Gonzalez, Robert Thai, Pauline Urquia, Anne-Cécile Van Baelen, Annalisa Meola, Ignacio Fernandez, Stéphanie Deville-Foillard, Ewan MacDonald, Léa Paolini, Frédéric Schmidt, Félix A. Rey, Michael S. Kay, Eric Tartour, Denis Servent, Ludger Johannes

**Author notes:** These authors contributed equally to this work (Anne Billet, Justine Hadjerci, Thi Tran, Pascal Kessler).

## Abstract

The success of mRNA-based vaccines during the Covid-19 pandemic has highlighted the value of this new platform for vaccine development against infectious disease. However, the CD8^+^ T cell response remains modest with mRNA vaccines, and these do not induce mucosal immunity, which would be needed to prevent viral spread in the healthy population. To address this drawback, we developed a dendritic cell targeting mucosal vaccination vector, the homopentameric STxB. Here, we describe the highly efficient chemical synthesis of the protein, and its *in vitro* folding. This straightforward preparation led to a synthetic delivery tool whose biophysical and intracellular trafficking characteristics were largely indistinguishable from recombinant STxB. The chemical approach allowed for the generation of new variants with bioorthogonal handles. Selected variants were chemically coupled to several types of antigens derived from the mucosal viruses SARS-CoV-2 and type 16 human papillomavirus. Upon intranasal administration in mice, mucosal immunity, including resident memory CD8^+^ T cells and IgA antibodies was induced against these antigens. Our study thereby identifies a novel synthetic antigen delivery tool for mucosal vaccination with an unmatched potential to respond to an urgent medical need.

## 1. Introduction

The COVID crisis has emphasized the need for efficient vaccines and revealed a place for mRNA formulations in this context. Current mRNA-based COVID vaccines effectively prevent severe forms of the disease but not infection [1,2]. These mRNA vaccines indeed do not (or only weakly) induce mucosal immunity in the ear-nose-throat area [3,4], which is required to avoid the transmission of the virus [5,6]. Only 3 nasally administered vaccines against influenza and COVID have been approved, all based on viral preparations [7,8]. Two of these encode SARS-CoV-2 spike protein and have entered the market only in China and India [8]. Of note, another virus-based vaccine expressing spike protein failed to induce significant mucosal responses [9]. Thereby, a continued need exists for new classes of mucosal vaccines. These should induce both IgA and mucosal CD8^+^ T cells, in particular resident memory CD8^+^ T lymphocytes (TRM) for which our teams and others have identified a major role in controlling infections and tumor growth in mucosal sites [10–14].

Shiga toxin B subunit (STxB) is a protein-based antigen delivery tool that targets dendritic cells. When conjugated to antigens, STxB induces cellular and humoral immunity against infectious disease or against tumors, leading to protective responses in peripheral tissues and lung and tongue mucosa [15–21]. STxB has several advantageous characteristics for its development as a vaccine vector: low intrinsic immunogenicity allowing for repeated vaccinations, high stability in circulation and ability to cross mucosal barriers (reviewed in [13]). STxB functions in synergy with adjuvants and immune checkpoint inhibitors by targeting different subpopulations of dendritic cells, and has no apparent toxicity. In previous vaccination studies, the homopentameric STxB was purified from bacteria which is time consuming, costly and plagued with endotoxin contamination [22]. Herein, we present a straightforward approach to obtain STxB via the synthetic route [23]: linear solid phase peptide synthesis of the 69 amino acid monomer sequence and highly efficient folding of the protein to obtain fully functional homopentamers without the need for liquid chromatography purification. We then produced a large number of STxB variants with functionalization sites at various positions of the protein. The most promising ones were assessed as fully synthetic vaccination vectors to evaluate their ability to induce TRM and IgA responses in the mucosa of the respiratory tract of mice.

## 2. Materials and methods

### 2.1. HPLC purifications and LC/ESI-MS analyses

#### 2.1.1. HPLC purifications of antigenic peptides

Purifications were done on Water Xbridge BEH 300 Prep C18 5 μm 30 × 150 mm columns with a Waters 2545 Quaternary Gradient Module, a Waters 2998 Photodiode Array Detector, and a Waters FlexInject. Solvents were: A - Milli-Q water; B - Acetonitrile. The flow rate was 30 mL/min. Pure fractions were freeze-dried using an ALPHA 2-4 LDplus freeze-dryer from CHRIST, with a Chemistry-HYBRID pump RC 6 from Vacuubrand.

#### 2.1.2. HPLC/ESI-MS analyses for synthetic and recombinant STxB

STxB samples were solubilized in guanidine hydrochloride (GdnHCl) (for crude peptides) or were directly injected in PBS (for folded or recombinant STxB) onto Waters XBridge BEH300 C18 3.5 μm 4.6 × 150 mm columns with an Agilent 1100 HPLC system coupled on-line to an ion-trap Bruker Esquire HCT mass spectrometer. Solvents were: A - 0.1 % TFA in Milli-Q water; B - 0.09 % TFA in acetonitrile. The flow rate was 0.8 mL/min. Gradients were: 5 min at 10 % B (or 10 min for Figure S10,S11 and S13) for accumulation at the head of the column, followed by a 20 min linear gradient from 10 % to 70 % B, then within 5 min from 70 % to 100 % B. Chromatogram was displayed at 214 nm. Mass spectra were acquired in Electron Spray Ionization (ESI) positive mode with high voltage capillary at 4500 V in ultra-scan mode (26000 m/z / sec scan) on an m/z 400-2000 scale and a target mass at m/z 1400. Smoothed multi-protonated spectra were deconvoluted into averaged mass spectra with Data Analysis software (Bruker-Daltonik, GmbH).

#### 2.1.3. UPLC/ESI-MS analyses for antigenic peptides and STxB-antigens conjugates

Samples were analysed with a Waters UPLC-MS comprised of an ACQUITY UPLC H-Class sample manager, an ACQUITY UPLC PDA eLambda Detector, and a Single Quadrupole Detector 2 for positive and negative ESI mass analyses. An ACQUITY UPLC BEH C18 1.7 μm 2.1 × 50 mm column was used. Solvents were: A - 0.1 % formic acid in Milli-Q water; B - 0.1 % formic acid in acetonitrile. The flow rate was 0.6 mL/min. Gradients were: 0.2 min 5 % B for accumulation at the head of the column, followed by 2.3 min linear gradient from 5 % to 95 % B. Antigenic peptide purities were calculated from surface peak integration at 214 nm from UPLC analyses.

### 2.2. STxB synthesis and folding

#### 2.2.1. Solid phase peptide synthesis

The solid phase synthesis of STxB was performed automatically on a Prelude or a Chorus instrument (Gyros Protein Technologies) in Fmoc/*t*Bu strategy. The following materials were purchased as indicated: Fmoc-Arg(Pbf)-Wang low-loading resin (0.29 theoretical loading) from Novabiochem for synthesis of acid C-terminal STxB, Rink Amide low-loading ChemMatrix resin (0.29 theoretical loading) from Sigma for synthesis of amide C-terminal STxB, amino acids and pseudoprolines from Novabiochem, Iris biotech or Gyros Protein Technologies, *N*-methylmorpholine (NMM), acetic anhydride (Ac_2_O), thioanisole, anisole, triisopropylsilane and acetic acid from Sigma Aldrich, HCTU (O-(1H-6-chlorobenzotriazole-1-yl)-1,1,3,3-tetramethyluronium hexafluorophosphate) from VWR, dichloromethane (DCM), diethyl ether, and piperidine from Carlo Erba, dimethylformamide (DMF) from Merck Millipore, *N*-Methyl-2-pyrrolidone (NMP) from BDH chemicals, trifluoroacetic acid (TFA) from Fisher Scientific. Pseudoprolines were incorporated regularly along the sequence and coupling optimizations were performed for some difficult positions highlighted in green (Figure 1b). Syntheses were performed on a 25 μmol scale for STxB syntheses in Table 1 and for C-terminal modified STxB variants presented in Table 2. A 12.5 μmol scale was used for other STxB azido-variants presented in Table S1.

**Table 1:** Synthesis and folding yields for STxB on two different synthesizers at a 25 μmol scale. The best yields were obtained with double coupling for 10 min and heating at 50 °C for difficult positions. Coupling at 70 °C did not yield further improvement (data not shown).

| Synthesis | Synthesizer | Use of pseudoprolines | Coupling conditions | Coupling conditions for difficult positions | Fmoc synthesis yield (%) | Crude peptide mass (mg) | Folding yield (%) | Overall yield (%) |
| --- | --- | --- | --- | --- | --- | --- | --- | --- |
| A | Prelude | No | 2x10 min RT | 2x10 min RT | 16.5 | 151.6 | $9 \pm 0.1$ | 1.5 |
| B | Prelude | Yes | 2x10 min RT | 2x10 min RT | 47.1 | 165.1 | $39 \pm 9.9$ | 18.4 |
| C | Prelude | Yes | 2x10 min RT | 4x10 min RT | 45.9 | 141.8 | $40.5 \pm 7.8$ | 18.6 |
| D | Chorus | Yes | 2x5 min RT | 2x5 min 50 °C | 19.4 | 147.8 | $15 \pm 1.4$ | 2.9 |
| E | Chorus | Yes | 2x10 min RT | 2x10 min 50 °C | 53.1 | 147.9 | $49 \pm 1.4$ | 26.0 |
| F | Chorus | Yes | 1x20 min RT | 1x20 min 50 °C | 49.9 | 158.6 | $39 \pm 8.5$ | 19.5 |

**Table 2:** Synthetic STxB variants with C-terminal modifications.

| Variant | Fmoc synthesis yield (%) | Crude peptide mass (mg) | Oxidation and folding yield (%) | Overall yield (%) | Retrograde trafficking |
| --- | --- | --- | --- | --- | --- |
| sSTxB | 47.1 | 165.1 | 39 | 18.4 | Functional |
| sSTxB(70C) | 42.8 | 111.3 | 28.4 | 12.2 | Functional |
| sSTxB(70AN<sub>3</sub>) | 47.7 | 120.9 | 17.5 | 8.3 | Functional |

**Figure 1:**
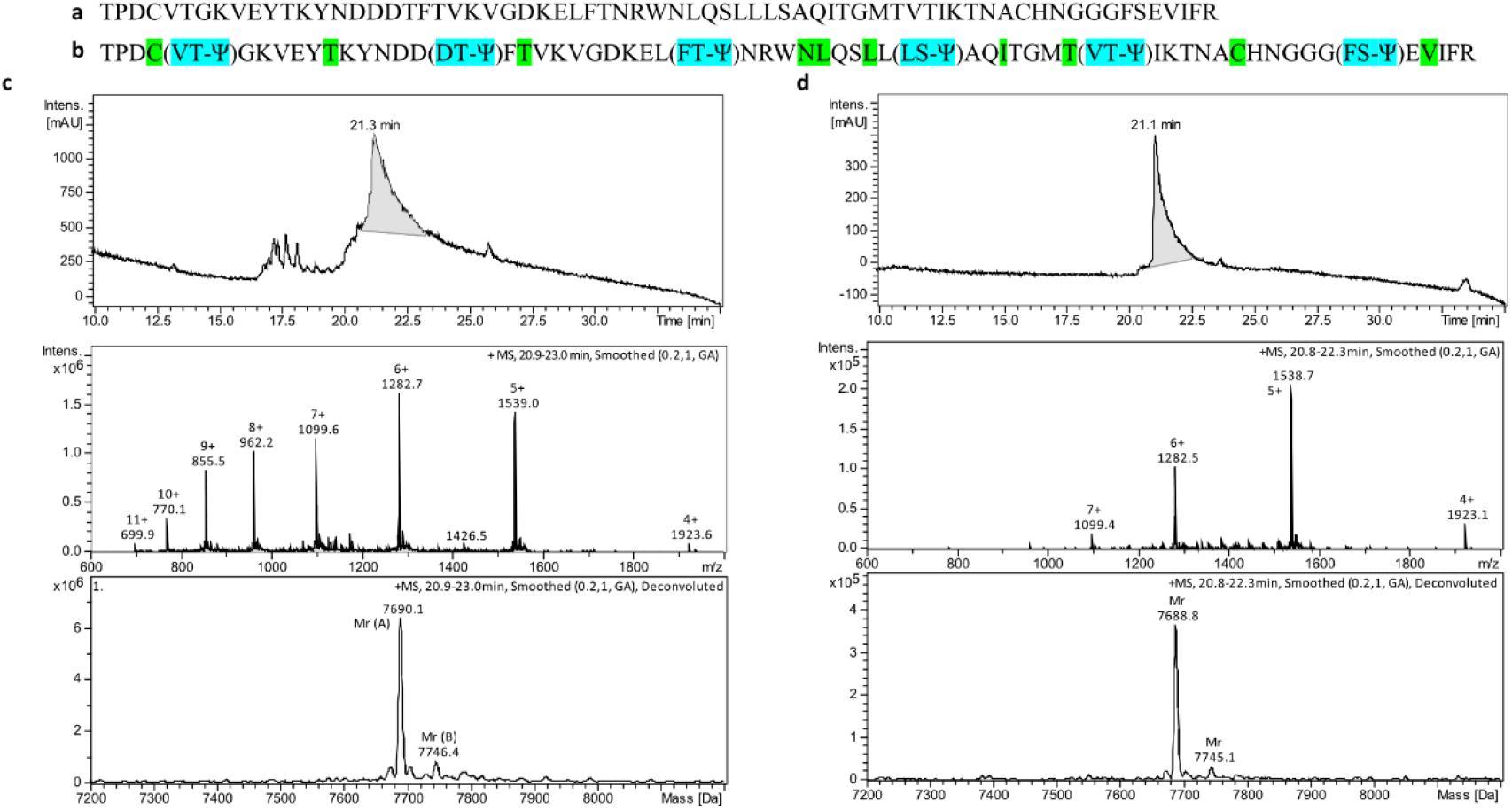
a) STxB SPPS sequence without optimization used for synthesis A, Table 1. All amino acids used were introduced with protected side chains. b) STxB SPPS sequence with optimizations. Pseudoprolines (highlighted in cyan) were inserted at different positions in the sequence to limit the aggregation of the peptide during elongation by solid phase peptide synthesis. Difficult positions for which coupling optimization were performed are highlighted in green. All amino acids were introduced with protected side chains. c) HPLC/ESI-MS graphs of newly synthesized STxB monomers showing the UV trace at 214 nm in the upper window, and multi-protonated positive ESI mass spectrum (middle window) which gives after deconvolution an average measured MW of 7690.1 ± 0.3 versus calculated MW= 7690.5. d) LC/ESI-MS graphs of refolded STxB homopentamers showing an average measured MW of 7688.8 ± 0.3 versus calculated MW= 7688.6.

Fmoc deprotection steps were done with 2 × 2 mL 2 min 20 % piperidine in NMP, followed by 3 × 3 mL 30 s NMP washes. Coupling steps were done for 5, 10 or 20 min with 1.3 mL 200 mM amino acid in NMP, 1 mL 250 mM HCTU in NMP, and 0.5 mL 1 M NMM in NMP, followed by 2 × 3 mL 30 s NMP washes. Coupling steps were performed once, or were repeated two to four times (as specified in Table 1). Systematic *N*-capping was done for 5 min with 2 mL 250 mM acetic anhydride in NMP and 0.5 mL 1 M NMM in NMP, followed by 3 × 3 mL 30 s NMP washes.

SPPS yield was determined from the last Fmoc deprotection step. Since capping was performed after each amino acid coupling, the amount of dibenzofulvene-piperidine adduct formed during piperidine-mediated deprotection of the last amino acid was directly related to the amount of full-length peptide present on the resin. This quantification was based on the dibenzofulvene-piperidine adduct absorbance at 301 nm. To do so, the deprotection solutions from the last amino acid were collected together and completed to 50 mL with DCM, before absorbance measurement at 301 nm. Yields were then calculated with the following formula (with a final volume of 50 mL and ε_fluorene_ = 7,800 M^-1^cm^-1^):

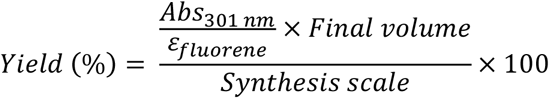

Resin cleavage was performed with 5 mL TFA/thioanisole/anisole/TIS/H_2_O (82.5/5/5/2.5/5) for 2 h under stirring. The cleavage solution was then precipitated in 40 mL cold diethyl ether. After 3 washes with cold diethyl ether, the precipitate was air dried and dissolved in aquous10 % acetic acid before freeze-drying. HPLC-MS analysis of STxB peptide is shown in Figure 1c.

#### 2.2.2. STxB oxidation and folding

Guanidine hydrochloride (GndHCl) was purchased from Calbiochem, sodium phosphate monobasic, sodium phosphate dibasic and dimethyl sulfoxide (DMSO) from Sigma-Aldrich.

##### In-solution oxidation with DMSO

Crude peptide was dissolved at 0.5 mg/mL in oxidation buffer (7 M GndHCl, 50 mM sodium phosphate, 2 % DMSO, pH adjusted to 8). The solution was incubated under stirring (220 rpm) for 24 h at 37 °C to form the disulfide bond (with holes in the lid to allow air oxidation). This oxidation method was used for all STxB variants described in the paper.

##### In-solution oxidation with disulfiram

Crude peptide was dissolved at 0.5 mg/mL in oxidation buffer (7 M GndHCl, 50 mM sodium phosphate, pH adjusted to 8, with 2 equivalents of disulfiram per equivalent of peptide). The solution was incubated under stirring (220 rpm) for 24 h at 37 °C to form the disulfide bond.

Ellman’s test procedure for the follow-up of oxidation in solution: 0.05 μmol of peptide (∼ 770 μL of oxidation solution at 0.5 mg/mL of STxB crude peptide) was diluted to 3 mL with 100 mM phosphate buffer pH 8, to which 100 μL of Ellman’s reagent solution (20 mg 5,5′-dithiobis(2-nitrobenzoic acid) (DTNB) in 10 mL 100 mM phosphate buffer pH 8) was added. The resulting solution was mixed and left to stand for 15 min. The absorbance of the solution was then measured at 410 nm after a blank with 100 μL Ellman’s reagent solution and 3 mL 100 mM phosphate buffer pH 8. Thiol concentration was calculated with the following formula calculation:

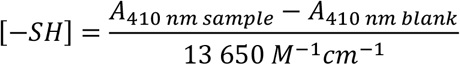

##### On-resin oxidation protocol

STxB was synthesized with STmp protecting groups on the two cysteines implicated in the intramolecular disulfide bond (Cys 4 and Cys 57), using the commercially available Fmoc-Cys(STmp)-OH amino acid. After SPPS, STmp was selectively removed on resin with 3 × 5 min treatments using 3 mL 5 % DTT, 0.1 M NMM in DMF (solution prepared with 1 g DTT, 220 μL NMM, and completed to 20 mL with DMF). The resin was washed 6 × 1 min with 3 mL DMF. On-resin oxidation was then performed under stirring for 30 min at room temperature with 3 mL DMF containing 2 eq *N*-chlorosuccinimide over free cysteines, thus corresponding to 4 eq over STxB monomer peptide. The resin was then washed 6 × 1 min with 3 mL DMF.

Few resin beads were collected before and after STmp deprotection and after oxidation to test for the presence of free thiols, using 2,2’-dithiobis-(5-nitropyridine) (DTNP). These resin beads were rinsed with ethanol, then incubated for 1-3 min with 200 μL DTNP solution (3 mg DTNP in 3 mL DMF). The apparition of a yellow color indicated the presence of free thiols (see Figure S9c).

The oxidized peptide was then cleaved from the resin using the standard cleavage protocol presented in the Solid phase peptide synthesis section. Oxidized and lyophilized peptide was dissolved at 0.5 mg/mL in 6 M GndHCl, 50 mM phosphate buffer pH 8 before folding dialyses.

##### Protein folding

After oxidation, folding occured at 4 °C upon dialyses against the following solutions with Slide-A-Lyzer™ dialysis cassettes 3500 MWCO from Thermo Scientific: 3 M GndHCl, 50 mM sodium phosphate pH 8.0, 5 mM EDTA for 6-10 h.

- 1 M GndHCl, 50 mM sodium phosphate pH 8.0, 1 mM EDTA overnight.
- PBS (0.14 M NaCl, 0.0027 M KCl, 0.01 M Phosphate buffer pH 7.4) for 2 times 4 h, then 1 time overnight.

After removal from the dialysis cassette, the solution was centrifuged for 20 min at 4 °C 4100 *g* to remove precipitated protein. The supernatant was concentrated at 4 °C, 2,000 *g* using Amicon Ultra Centrifugal filters 10,000 MWCO from Merck Millipore. After concentration, the supernatant was centrifuged again at 17,000 *g*, 20 min at 4 °C to remove precipitated protein. Protein concentration in the remaining supernatant was determined by measuring absorbance at 280 nm with baseline correction at 340 nm using a Nanodrop 2000c (Thermo Scientific). The Gill and von Hippel molar extinction coefficient [24] was used to calculate the concentration of STxB conjugates (ε = 41,850 M^-1^cm^-1^). Except when mentioned otherwise, concentrations in this paper correspond to STxB pentamers. Small aliquots were flash-frozen in liquid nitrogen and stored at -20 °C.

STxB nomenclature: sSTxB or rSTxB for synthetic or recombinant STxB, respectively, followed by the original amino acid in the STxB sequence, its position, and then the new amino acid. For C-terminal modifications, position 70 is indicated followed by the additional amino acid. SPPS yields for sSTxB are detailed in Table 1 in function of coupling conditions. HPLC/ESI-MS graphs are presented in Figure 1 and in Figures S1-7 (MS m/z C_339_H_528_N_90_O_108_S_3_ Average MW calculated: 7688.6, found: between 7688.2 and 7688.6; [M+4H^+^]^4+^ calculated: 1923.1, found: 1923.0, [M+5H^+^]^5+^ calculated: 1538.7, found: between 1538.6 and 1538.7; [M+6H^+^]^6+^ calculated: 1282.4, found: between 1282.4 and 1282.5; [M+7H^+^]^7+^ calculated: 1099.4, found: between 1099.3 and 1099.4; [M+8H^+^]^8+^ calculated: 962.1, found: between 962.1 and 962.3). In comparison, HPLC analyse of rSTxB is presented Figure S8. sSTxB(70C) and sSTxB(70AN_3_) SPPS yields are detailed in Table 2 and HPLC/ESI-MS graphs are presented in Figures S10-13. Characterizations are detailed in Supplementary data. Other STxB-azido variants SPPS yields are presented in Table S1.

### 2.3. Biophysical characterization of STxB

#### 2.3.1. Circular dichroism

Circular dichroism experiments were performed on a J-815 CD JASCO instrument, using a 1 mm quartz cell. STxB samples at 6 μM concentration in PBS were used. Far-UV (190-250 nm) circular dichroism spectra were recorded at 0.1 nm resolution at 20 °C with a scanning speed of 50 nm/min and 1 s response time. Background (PBS spectra) was subtracted from raw data. Three scans were acquired and averaged for each sample. The molar ellipticity was calculated from the circular dichroism intensity in mdeg.

#### 2.3.2. Detection of DNA contamination at 260 nm and scattering at 340 nm

Protein quantification at 280 nm was carried out by recording a full spectrum between 240 and 340 nm. Measurements were done with 60 μL of PBS and sample at 20 °C in a 1 cm quartz cell, reference 105.202-QS.10 (Hellma), using a JASCO V-750 spectrophotometer. A baseline subtraction at 340 nm was performed with the Spectragryph software (https://www.effemm2.de/index.html) to accurately calculate the protein concentration. The cuvettes were cleaned with 2 % Hellmanex/70 % ethanol/water.

#### 2.3.3. Assessment of the sample homogeneity and quantification

Dynamic light scattering analysis was performed on a DynaPro Plate Reader III (Wyatt). A volume of 20 μL of sample at 0.1 mg/mL was loaded in a 384-well microplate (Corning ref 3540), with 10 acquisitions of 5 s each at 20 °C, monitored with the DYNAMICS version V7.10.0.21 software (Wyatt). Measurements were performed in triplicate.

#### 2.3.4. Thermal denaturation by nanodifferential scanning fluorimetry (nanoDSF)

Thermal denaturation was performed on the Prometheus NT.48 instrument (Nanotemper) from 20 to 95 °C with a 1 °C/min heating rate. Tryptophan fluorescence emission was monitored at 330 nm and 350 nm as a function of increasing temperature. The capillaries were filled with 10 μl of protein at 0.1 mg/mL and the intrinsic fluorescence signal expressed as the 350 nm/330 nm emission ratio was plotted as a function of temperature. Measurements were performed in triplicate.

### 2.4. Cell culture

HeLa cells were cultured at 37 °C under 5 % CO_2_ in Dulbecco’s modified Eagle’s medium (DMEM, Invitrogen), supplemented with 10 % fetal bovine serum, 0.01 % penicillin-streptomycin, 4 mM glutamine and 5 mM pyruvate (complete medium).

### 2.5. Intracellular trafficking assay

The day before the experiment, cells were seeded on glass lamellae in 4-well dishes, 80,000 cells/well. On the day of the experiment, cells were incubated for 30 min at 4 °C in 500 μL of 40 nM STxB in ice-cold complete medium for binding, followed by 3 washes with PBS^++^ (PBS, 0.5 mM MgCl_2_, 1 mM CaCl_2_). Complete medium at 37 °C was added to cells, which were then incubated for 50 min at 37 °C for synchronized internalization. Cells were washed 3 times with PBS^++^, fixed with 4 % paraformaldehyde (PFA) in PBS for 15 min, washed once with 50 mM NH_4_Cl, and incubated with 50 mM NH_4_Cl for 30 min to quench the PFA. Cells were washed 3 times with PBS/BSA/saponin (PBS/0.2 % w/v bovine serum albumin/0.02 % w/v saponin) and permeabilized at room temperature for 30 min in PBS/BSA/saponin. Lamellae were incubated on 30 μL of antibody dilution into PBS/BSA/saponin for 30 min at room temperature, then washed 3 times with PBS/BSA/saponin. Primary antibodies were a homemade mouse monoclonal anti-STxB antibody (13C4, used at 1/250 dilution), and a homemade rabbit polyclonal antibody against the Golgi marker giantin (used at 1/100 dilution). The secondary antibodies were Cy3-coupled anti-mouse and Alexa488-coupled anti-rabbit IgGs, used at 1/100 dilution each. Lamellae were washed in water and then deposited onto slides with 6 μL of Fluoromont G with Hoechst (2.5 μg/mL). Polymerization was performed for 30 min at 37 °C.

### 2.6. Microscopy

#### 2.6.1. Retrograde trafficking

Slides were observed on microscopes of the Cell and Tissue Imaging platform (PICT-IBiSA) at Institut Curie:

- Upright Leica DM6000 microscope with a CCD 1392×1040 CoolSnap HQ2 (pixel size: 6.45 μm) camera from Photometrics with illumination by a Lumen 200 lamp from Prior Scientific and a 63x HCX Plan Apo objective.
- Upright Leica DM6B microscope with a sCMOS Orca Flash 4.0 V2 camera (pixel size: 6.5 μm) from Hamamatsu with illumination by Metal-Halide EL 6000 lamp from Leica and a 100x HCX PL APO objective.

#### 2.6.2. Colocalization

Images were captured using the inverted Eclipse Ti-E (Nikon) with spinning disk CSU-X1 (Yokogawa) and 60x CFI Plan Apo. Stacks of 16 images at 0.2 μm depth were integrated with Metamorph software by Gataca Systems. Single cells were analysed using the colocalization threshold tool in Fiji ImageJ software (National Institutes of Health) [25].

### 2.7. Preparation of STxB-antigen conjugates

For each conjugation, the concentration of recombinant or synthetic STxB was between 40 and 50 μM. The concentration of STxB conjugates was determined by measuring absorbance at 280 nm with baseline correction at 340 nm using a Nanodrop 2000c (Thermo Scientific). Gill and von Hippel molar extinction coefficients [24] were used to calculate the concentration of STxB conjugates (ε = 41,850 M^-1^cm^-1^ for STxB-SL8 and STxB-G15F conjugates, and ε = 75,700 M^-1^cm^-1^ for STxB-RBD conjugates).

#### 2.7.1. RBD production

The receptor binding domain (RBD; residues 331-528) from SARS-CoV-2 was cloned in a vector suitable for expression in Schneider’s Drosophila Line 2 cells (ATTC CRL-1963). In detail, a synthetic codon-optimized gene was cloned into a modified pMT/BiP plasmid resulting in the translation of the protein in frame with an enterokinase cleavage site and double strep-tag at the C-terminus. Stable clones were generated as previously reported [26]. Cells were amplified and protein production was induced by adding 4 μM CdCl_2_. After 6 days of incubation, cell supernatant was collected, centrifuged and purified on a Streptactin column. The eluted fractions were pooled and run on a Superdex 75 column equilibrated with 10 mM Tris-HCl (pH 8.0), 100 mM NaCl. The peak corresponding to monomeric protein was harvested, quantified and incubated with enterokinase (Biolabs) overnight at 4 °C in the presence of 2 mM CaCl_2_. To separate cleaved from uncleaved protein, the mixture was then purified on a Streptactin column, the flow through was collected and run on Superdex 75 in 10 mM Tris-HCl (pH 8.0), 100 mM NaCl. The peak corresponding to the monomeric, untagged protein was collected, concentrated, quantified and kept at -80 °C until further use.

#### 2.7.2. Conjugation between sSTxB(70C) and RBD

The RBD protein was incubated for 30 min at 21 °C and 750 rpm with a 10 molar equivalent of sulfo-m-maleimidobenzoyl-*N*-hydoxysuccinimide ester (Sulfo-MBS) (Thermo Scientific; MW 416.30). The excess crosslinker was removed by passing the protein through a PD-10 column using a PBS, 10 mM EDTA buffer. sSTxB(70C) was mixed with one molar equivalent of the activated RBD protein. The conjugation reaction was performed overnight at 21 °C and 750 rpm. The formation of sSTxB(70C)-RBD conjugates was verified by western blotting using rabbit anti-STxB polyclonal antibody (produced upon request by Covalab) and rabbit anti-SARS-CoV-2 spike protein (RBD) polyclonal antibody (Thermo Fisher Scientific; catalog # PA5-114451).

#### 2.7.3. Conjugation between recombinant and synthetic STxB(70C) and bromoacetyl-SL8 or bromoacetyl-G15F

Recombinant rSTxB(70C) was produced as previously described [27]. Recombinant and synthetic STxB(70C) were dialyzed against 50 mM borate buffer, 150 mM NaCl pH 9. The conjugation reaction was performed overnight at 21 °C and 750 rpm with 3 equivalents of Br-CH_2_CO-OVA_257-264_ peptide (SL8) or Br-CH_2_CO-E7_43-57_ peptide (G15F) (both produced by Polypeptide Group) per STxB monomer with 10 % DMSO to keep the peptides soluble. Of note, three additional amino acids were added at the N-terminus of OVA_257-264_ peptide to enable efficient processing of the antigen after conjugation to STxB (leading to the sequence QLESIINFEKL). Conjugates were purified from excess of free antigenic peptides by several dialysis steps at 4 °C against PBS using Slide-A-Lyzer 20,000 MWCO dialysis cassettes (Thermo Scientific). STxB(70C)-SL8 and STxB(70C)-G15F conjugate formation and the absence of remaining free SL8 or G15F after purification were validated by UPLC-MS.

#### 2.7.4. DBCO-SL8 and DBCO-PEG4-SL8 syntheses

Cys-SL8 (H-CQLESIINFEKL-OH) was synthesized on a Prelude instrument at a 25 μmol scale using the conditions described in the Solid phase peptide synthesis section, with 2 coupling repetitions of 10 min per amino acid. 10 mg crude was reacted with 1.1 equivalent of commercial DBCO-Mal (Iris biotech; CAS: 1395786-30-7; MW 427.45 g/mol) or DBCO-PEG4-Mal (Iris biotech; CAS: 1480516-75-3; MW 674.74 g/mol) in 4 mL 40 % acetonitrile, 60 % 50 mM ammonium bicarbonate buffer pH 7 for 1 h, prior to freeze-drying. The products were purified by HPLC with a 30 min gradient from 5 to 100 % acetonitrile in water yielding 2.1 mg DBCO-SL8 in powder form (yield 16 %, purity 93 %, MS m/z C_89_H_126_N_18_O_24_S [M+H^+^]^+^ calculated: 1865.15, found: 1864.7; [M+2H^+^]^2+^ calculated: 933.1, found: 932.9; [M-H^-^]^-^ calculated: 1863.15, found: 1863.2) and 2.5 mg DBCO-PEG4-SL8 (yield 17 %, purity 78 %, MS m/z C_100_H_147_N_19_O_29_S [M+H^+^]^+^ calculated: 2112.4, found: 2112.4; [M+2H^+^]^2+^ calculated: 1056.7, found: 1056.6; [M+3H^+^]^3+^ calculated: 704.8, found: 704.8; [M-H^-^]^-^ calculated: 2110.4, found: 2110.9).

#### 2.7.5. Conjugation between sSTxB(70AN_3_) and DBCO-SL8 or DBCO-PEG4-SL8

sSTxB(70AN_3_) was dialyzed for 4 h against 50 mM borate buffer, 150 mM NaCl pH 9 using a Slide-A-Lyzer 0.5 - 3 mL 10,000 MWCO dialysis cassette (Thermo Scientific). The conjugation reaction was performed overnight at 21 °C and 750 rpm with 2 equivalents of DBCO-SL8 or DBCO-PEG4-SL8 per STxB monomer and 10 % DMSO to keep the peptides soluble. The excess of free SL8 peptides was removed on Zeba™ Spin Desalting Columns, 7,000 MWCO, 2 mL (Thermo Scientific), followed by an overnight dialysis step against PBS using Slide-A-Lyzer 0.5 - 3 mL 10,000 MWCO dialysis cassettes (Thermo Scientific). sSTxB(70AN_3_)-SL8 conjugates formation and the absence of remaining free SL8 after purification were validated by UPLC-MS. Characterization of conjugates is detailed in Supplementary data.

#### 2.7.6. DBCO-G15F and DBCO-PEG4-G15F syntheses

Cys-G15F (H-CGQAEPDRAHYNIVTF-OH) was synthesized on a Chorus instrument at a 25 μmol scale using the conditions described in the Solid phase peptide synthesis section, with 2 coupling repetitions of 10 min per amino acid. 20 mg crude was reacted for 1 h with 1.2 equivalent of commercial DBCO-Mal (Iris biotech; CAS: 1395786-30-7; MW 427.45 g/mol) or DBCO-PEG4-Mal (Iris biotech; CAS: 1480516-75-3; MW 674.74 g/mol) in 6 mL 40 % acetonitrile, 60 % 50 mM ammonium bicarbonate buffer pH 7, prior to freeze-drying. The products were purified by HPLC with a 25 min gradient from 10 to 70 % acetonitrile in water yielding in powder form 7.8 mg DBCO-G15F (yield 31.5 %, purity 85 %, MS m/z C_104_H_138_N_26_O_29_S [M+H^+^]^+^ calculated: 2249.4, found: 2248.6; [M+2H^+^]^2+^ calculated: 1125.2, found: 1125.2; [M+3H^+^]^3+^ calculated: 750.5, found: 750.5; [M-H^-^]^-^ calculated: 2247.4, found: 2246.7) and 6.4 mg DBCO-PEG4-G15F (yield 23.3 %, purity 83 %, MS m/z C_115_H_159_N_27_O_34_S [M+H^+^]^+^ calculated: 2496.7, found: 2493.9; [M+2H^+^]^2+^ calculated: 1248.9, found: 1248.9; [M+3H^+^]^3+^ calculated: 832.9, found: 832.7; [M-H^-^]^-^ calculated: 2494.7, found: 2493.8).

#### 2.7.7. Conjugation between sSTxB(70AN_3_) and DBCO-G15F or DBCO-PEG4-G15F

The conjugation reaction was performed overnight at 21 °C and 750 rpm with 3 equivalents of DBCO-G15F or DBCO-PEG4-G15F per STxB monomer and 10 % DMSO to keep the peptides soluble. The excess of free G15F peptide was removed by overnight dialysis against PBS using Slide-A-Lyzer 0.5 - 3 mL 20,000 MWCO dialysis cassettes (Thermo Scientific). The formation of sSTxB(70AN_3_)-G15F conjugates and the absence of remaining free G15F peptides after dialysis were validated by UPLC-MS. Characterization of conjugates is detailed in Supplementary data.

### 2.8. In vivo crosspriming

#### 2.8.1. Mice and vaccination

Female BALB/cJ and C57bl/6J mice were purchased from Janvier Labs and used at 8-10 weeks of age. All mice were housed in INSERM U970-PARCC animal facility under specific pathogen-free conditions. Experimental protocols were approved by Université Paris Cité ethical committee (approval n°29315) in accordance with European guidelines (EC2010/63). Anesthetized mice were immunized twice at day 0 and day 14 by intranasal vaccination with 0.5 nmol of various vaccines associated with alpha-galactosylceramide (α-GalCer) (Funakoshi, Tebu-bio) or c-di-GMP (InvivoGen) as an adjuvant.

On day 21, mice were sacrificed, bronchoalveolar lumen fluid (BAL) was obtained by flushing the lungs with PBS-EDTA 0.5 mM via a cannula inserted in the trachea (5 washes x 1 mL).

Lungs were perfused with PBS-EDTA 0.5 mM, and digested in RPMI-1640 medium supplemented with 1 mg/mL collagenase type IV (Life Technologies/Thermofisher) and 30 μg/mL DNase I (Roche). Lung cells were dissociated using the GentleMACS (Miltenyi Biotec) lung programs 1 and 2, with gentle shaking for 30 min at 37 °C in between both steps. Then, the obtained single-cell suspensions were filtered through a 70-μm strainer, washed with PBS-FBS 2 %, suspended in 40 % Percoll solution, layered over 75 % Percoll solution (Sigma-Aldrich), and centrifuged for 20 min at RT and 600 *g*. Interface cells were collected and washed.

#### 2.8.2. Flow cytometry

After FcR blocking with CD16/32 Ab (clone 93, ebioscience/Life Technologies), cells were first incubated for 30 min at room temperature with PE-conjugated K^b^OVA_257-264_ or D^b^E7_49–57_ tetramers (Immudex, Bredevej 2A, 2830 Virum). Cells were then washed and stained for 20 min at 4 °C with the following antibodies: anti-mouse CD8α APC-efluo780 (clone 53-6-7, ebioscience/ Thermofisher), anti-mouse CD8β AF700 or BUV495 (clone YTS156, eBioscience), CD3 PercpCy5.5 (clone 145 2C11, ebioscience/Life Technologies), CD103 Pacific Blue (clone 2E7, Biolegend), and CD49a APC (Miltenyi Biotec). Acquisitions were performed on BD Fortessa X20 (Becton Dickinson), and data were analyzed on live single cells with FlowJo Software (BD), as recently described [28].

#### 2.8.3. Ex vivo ELISpot assays

CD8^+^ T lymphocytes were enriched from lung cell suspensions prepared according to the manufacturer’s instructions with the CD8^+^ T cell isolation kit (Stem Cell Technologies). Purified CD8^+^ T cells were incubated in ELISpot plates in the presence of medium or a pool of S1 peptides (1 μg/mL) covering the spike receptor binding domain (RBD) sequence of SARS-CoV-2 (Wuhan strain), or a pool of S2 peptides (1 μg/mL) derived from spike outside the RBD sequence. These S1 and S2 pools were purchased from JPT peptide technologies.

ELISpot plates were incubated for 18 to 21 hours at 37 °C with 5 % CO_2_. The IFNγ spots were revealed following the manufacturer’s instructions (Gen-Probe Diaclone). Spot-forming cells were counted with the C.T.L. Immunospot system (Cellular Technology Ltd.).

#### 2.8.4. Measurement of RBD-specific IgA and IgG

Recombinant SARS-CoV-2 spike RBD protein (R&D Biotechne,#10534-CV-100) in sodium carbonate buffer pH 9.6 was coated at 2 μg/mL (75 μL/well) overnight at 4 °C onto 96-well Maxisorp clear plates. All the following steps were done at room temperature. The coating buffer was removed, plates were washed 3 times with PBS-Tween 20 0.05 %, and blocked for 1 h in PBS-BSA 1 % (200 μL/well). 50 μL of diluted BAL were added and incubated for 90 min with gentle shaking. Plates were washed 3 times with PBS-Tween 20 0.05 %, followed by the addition of 1/10,000 goat anti-mouse IgA-biotin (Southern Biotech, #1040-08) or 1/10,000 goat anti-IgG-biotin (Southern Biotech #1030-08) diluted in PBS-BSA 1 % (100 μL/well). After incubation for 1 h, plates were washed 4 times with PBS-Tween 20 0.05 %, and 1/5,000 HRP-Steptavidin (100 μL/well) was added for 1 h (Southern Biotech, #7105-05). Plates were washed 4 times again with PBS-Tween 20 0.05 %, and 50 μL of 3,3’,5,5’-Tetramethylbenzidine (TMB) substrate (Eurobio Scientific, # 5120-0050) was added. After 10 min of incubation, reactions were stopped with 50 μL of TMB stop solution (KPL, #50-85-05). The absorbance at 450 nm was immediately measured on a Spectrostar microplate reader (BMG Labtech).

### 2.9. Data analysis and figures

Prism software (GraphPad) was used for statistical analysis and graph plotting, Fiji ImageJ software (National Institutes of Health) [25] for microscopy, Chemdraw software to design molecules and reaction schemes, and Affinity Publisher software to draw figures.

## 3. Results and discussion

### 3.1. Linear full-length STxB synthesis

STxB monomer syntheses were performed at room temperature or 50 °C, on automated Prelude or Chorus peptide synthesizers, respectively. In room temperature conditions, Fmoc synthesis and folding yields were 16.5 % and 9 %, respectively, leading to a total yield of only 1.5 % (Table 1, Synthesis A with the sequence presented Figure 1a). In a first step of optimization, pseudoprolines were regularly incorporated along the sequence to diminish peptide chain aggregation during elongation (Figure 1b). For positions known to be difficult for coupling, coupling temperatures (room temperature, 50 °C or 70 °C), time (5, 10 or 20 min) and the number of couplings (between 1 and 4) were also optimized. The best overall yield (26%) was obtained with 2 × 10 min couplings at RT or 50 °C for difficult positions (Table 1, Synthesis E) and is over 17-fold compared with STxB synthesis without optimization (Table 1, Synthesis A).

The crude peptide obtained after the TFA cleavage corresponded mostly to full-length STxB monomer, with only a few truncated products (Figure 1c).

Each monomer of STxB contains an intramolecular disulfide bond. We explored different conditions for its oxidation before folding. In a first approach, synthesized monomers were dissolved at 0.5 mg/mL in 7 M guanidine hydrochloride (GndHCl) solution at a slightly alkaline pH (50 mM phosphate pH 8) in the presence of atmospheric oxygen or oxidizing compounds (DMSO or disulfiram) [29,30]. Oxidation was followed by absorbance using Ellman’s reagent (DTNB) (Figure S9a). Disulfiram effectively accelerated the reaction, with half times of 1 h, whereas the addition of 2 % DMSO had no effect over air oxidation alone with half times of 4 h. In an alternative approach, oxidation was directly performed on the synthesis resin. For that, we used cysteines with the trimethoxyphenylthio (STmp) protecting group, which after synthesis was selectively removed with 5 % DTT. Then, *N*-chlorosuccinimide (NCS) was used for on-resin oxidation, which was complete after 30 min (Figure S9b), as shown by free thiol detection using 2,2’-dithiobis-(5-nitropyridine) (DNTP) (Figure S9c). The on-resin oxidation strategy took in total only 1 h and was thereby much faster than in-solution oxidation.

For folding of oxidized STxB monomers into functional homopentamers, the former were solubilized in 7 M GndHCl in phosphate buffer pH 8. GndHCl was then removed by sequential dialyses, reducing its concentration in a stepwise manner from 7 M to 0 M in PBS. Of note, this dialysis scheme concomitantly allowed to remove the oxidizing agent and peptides that had failed to fold correctly and that therefore precipitated. The homogeneity that was obtained after dialysis and centrifugation was such that no further purification was needed (Figure 1d and Figure S1-6). Furthermore, the contaminant glutathione adducts that are found on recombinant STxB purified from bacteria were absent altogether. In contrast, *t*-butyl adduct traces were detected in synthetic STxB. Assuming that *t*-butyl-derivatives of STxB had the same electron spray ionization behaviour as fully deprotected STxB, *t*-butyl-STxB represented 16% and 8% of reduced or refolded STxB monomers, respectively (Figures 1c-d). We therefore tested whether the conformational stability and functionality of synthetic STxB were preserved despite these adducts.

### 3.2. Synthetic STxB is very similar to recombinant STxB

The biophysical properties of recombinant and synthetic STxB homopentamers were compared. Circular dichroism analysis revealed very similar secondary structures for the two proteins, demonstrating that the synthetic protein folded correctly (Figure 2a). The size distribution was determined by dynamic light scattering (DLS), revealing that both proteins had hydrodynamic radii between 2.5 and 2.9 nm in solution (Figure 2b, c), corresponding to the expected size of STxB [31]. Two other populations with radii between 10 and 100 nm (peaks 2 and 3) were present at very low mass percentages and likely corresponded to soluble aggregates (Figure 2b, c). The most striking difference was a 4^th^ peak of 1.4 % in mass that was present only for synthetic STxB (Figure 2c). With a radius of around 1 μm, this material likely corresponded to non-soluble aggregates. Finally, thermal denaturation was monitored by nanodifferential scanning fluorimetry (nanoDSF). A small difference of 1.1 °C for the melting temperatures (Tm) of both proteins was measured (Figure 2d). Taken together, the biophysical properties of recombinant and synthetic STxB were very similar.

**Figure 2:**
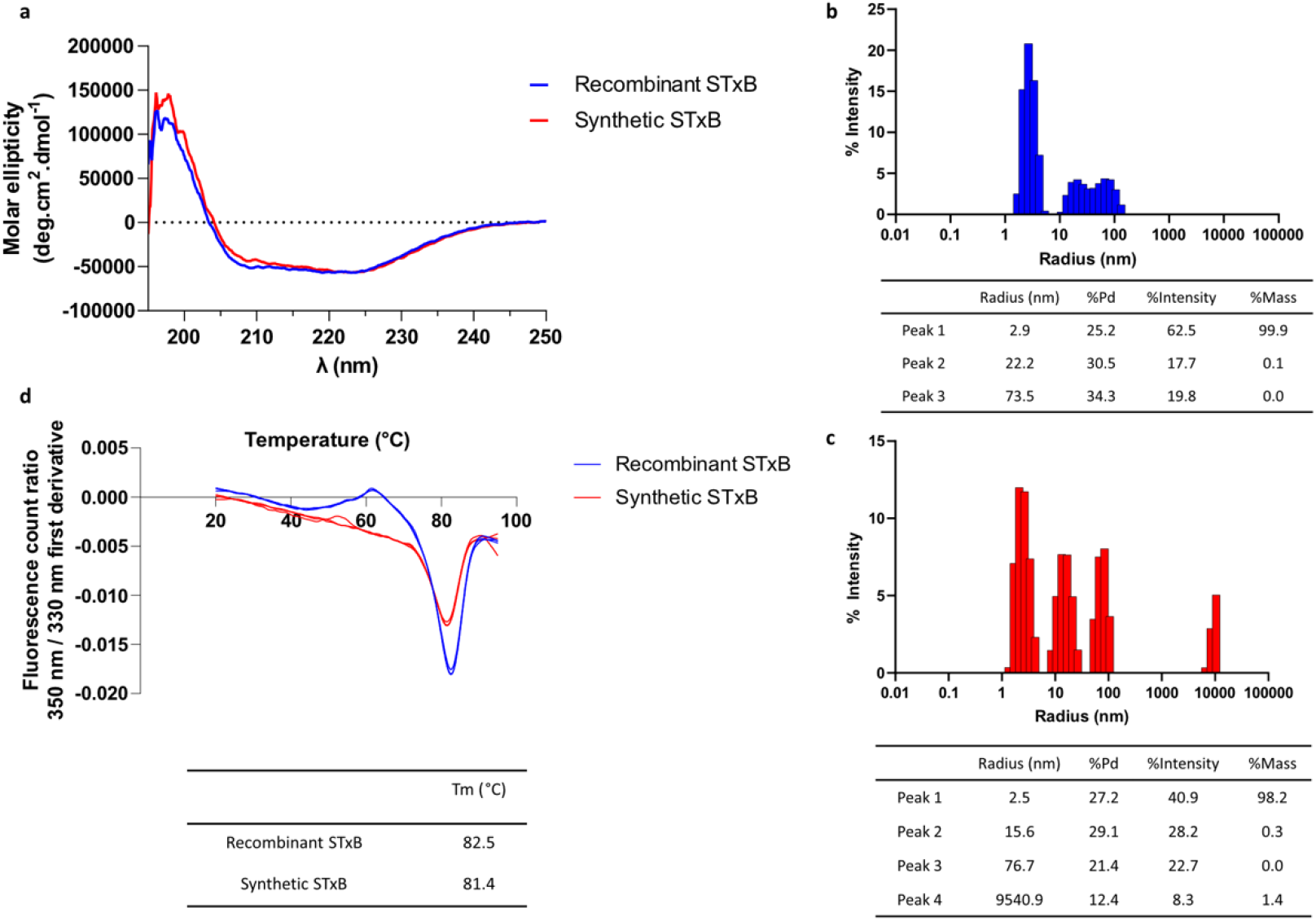
Biophysical analysis of recombinant and synthetic STxB. a) Circular dichroism revealed that recombinant and synthetic STxB had similar secondary structure contents. b) Analysis of the heterogeneity of recombinant STxB in PBS. The major population represented pentameric STxB with a radius of 2.9 nm. The other populations between 10 and 100 nm were attributed to soluble aggregates of STxB. c) Analysis of the heterogeneity of synthetic STxB in PBS. The behavior of the synthetic STxB was similar to that of recombinant STxB with the main population at 2.5 nm which represented more than 98% in mass. The other populations were aggregates. d) Thermal denaturation of STxB was measured by nanoDSF. The analysis of the first derivatives of fluorescence count ratios at 350 nm/330 nm in function of temperature revealed that the melting temperatures (Tm) of recombinant and synthetic STxB were very similar.

A key characteristic of STxB is its capacity to undergo retrograde trafficking from the plasma membrane to the Golgi apparatus, which depends entirely on the binding to its cellular receptor, the glycosphingolipid Gb3 [32]. To test the functionality of synthetic STxB, we compared its retrograde trafficking to that of recombinant STxB. For this, Gb3-expressing HeLa cells were incubated at 37 °C with recombinant or synthetic STxB. Cells were then fixed and labeled for STxB and the Golgi marker giantin. Both types of STxB proteins were detected in a perinuclear location (Figure 3a) where they colocalized to the same extent with giantin (Figure 3b). Very clearly, both synthetic and recombinant STxB were indistinguishable in their capacity to undergo retrograde trafficking to the Golgi.

**Figure 3:**
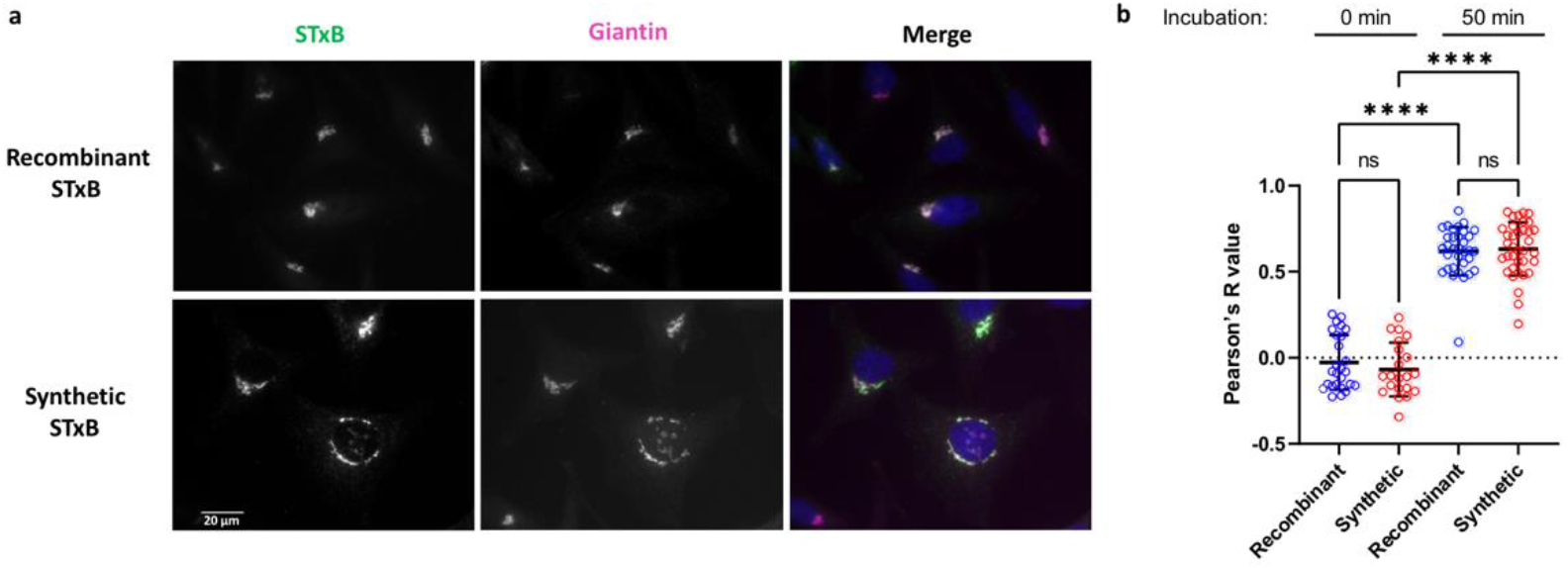
Retrograde trafficking of synthetic and recombinant STxB in HeLa cells. 40 nM STxB was bound to cells for 30 min at 4 °C, followed by PBS washes and incubation for 50 min at 37 °C for synchronized internalization. a) Immunofluorescence images of recombinant and synthetic STxB. Merge: STxB in green, Golgi in magenta, and DNA dye Hoechst in blue. Scale bars: 20 μm. b) Quantification of colocalization of STxB with giantin in the indicated conditions and at the indicated time points. ****P < 0.0001, ns: non-significant.

### 3.3. Chemical synthesis opens new possibilities for bioorthogonal conjugations with STxB

The capacity to obtain STxB in a highly efficient linear chemical synthesis scheme opened the possibility of producing STxB variants with defined functionalization sites. Two types of modifications were chosen: unnatural azide-containing amino acids for conjugation via copper-free click chemistry, and the addition of a cysteine on the C-terminus for coupling via sulfhydryl-specific chemistries. Positions for unnatural azide-containing amino acid substitutions were selected based on the analysis of the STxB structure in complex with an analogue of its receptor (Protein Data Bank: 1BOS). Key criteria were exposure at the surface of the homopentamer and non-engagement in Gb3 binding sites.

17 azido variants were synthesized with overall yields between 7 and 24 % (detailed in Table S1). For the majority of these, azidolysine (KN_3_) was used. For two positions located close to the membrane interaction surface of STxB, a shorter azidoalanine (AN_3_) was preferred to reduce the risk of interfering with receptor binding. At position 11, 4-azidomethylphenylalanine was used as it was structurally most closely related to the natural tyrosine. Their functionality in retrograde trafficking were also tested (Table S1). Overall, the incorporation of azide-containing amino acids turned out to be feasible at most positions chosen of the STxB sequence, which again documented a high degree of structural robustness of the protein. For C-terminal modifications involving the addition of a natural cysteine (termed sSTxB(70C)) or of non-natural azidoalanine (termed sSTxB(70AN_3_)), total yields were reduced compared to wild-type STxB. (Table 2). However, both STxB variants had excellent purity after folding (Figure S10-13) and were fully functional (Figure S14).

### 3.4. Synthetic STxB: an efficient delivery tool for mucosal vaccination

To address the potential clinical interest of synthetic STxB as an antigen delivery vector, sSTxB(70C) and sSTxB(70AN_3_) were chemically coupled to different types of antigens. The first one was the receptor binding domain (RBD) of the SARS-CoV-2 spike protein (Figure 4a for the chemical coupling scheme, and Figure S15 for conjugate analyses), which is required for virus entry into target cells by binding to the ACE2 receptor [33]. After intranasal immunization, sSTxB(70C)-RBD induced functional CD8^+^ T cells against RBD in the lung, as detected by Elispot with a pool of peptide that covered the entire S1 domain, including RBD (Figure 4b). With sSTxB(70C)-RBD, the frequency of IFNγ-producing CD8^+^ T cells was 4 times higher than that observed after mucosal vaccination with non-vectorized RBD (Figure 4b). No response was detected when the pool of S1 peptides was replaced with a pool of S2 domain-derived peptides that did not include RBD, reinforcing the specificity of the cellular response (data not shown). Importantly, sSTxB(70C)-RBD also induced 7 and 8-times higher concentrations of, respectively, IgA and IgG in bronchoalveolar fluid than non-vectorized RBD (Figure 4c-d). To benchmark synthetic STxB as an antigen delivery vector, we compared its immunomodulation efficiency to that of recombinant rSTxB(70C) that was used in previous anticancer vaccination studies with antigens from a model protein, i.e., the SL8 peptide from ovalbumin [34], and the G15F peptide from the E7 protein of HPV16 [17,20,21,35]. The chemical coupling schemes for sSTxB(70C)-SL8, sSTxB(70C)-G15F, sSTxB(70AN_3_)-SL8 and sSTxB(70AN_3_)-G15F are shown in Figure 5a, and their chemical analyses and retrograde trafficking in Figures S16-19.

**Figure 4:**
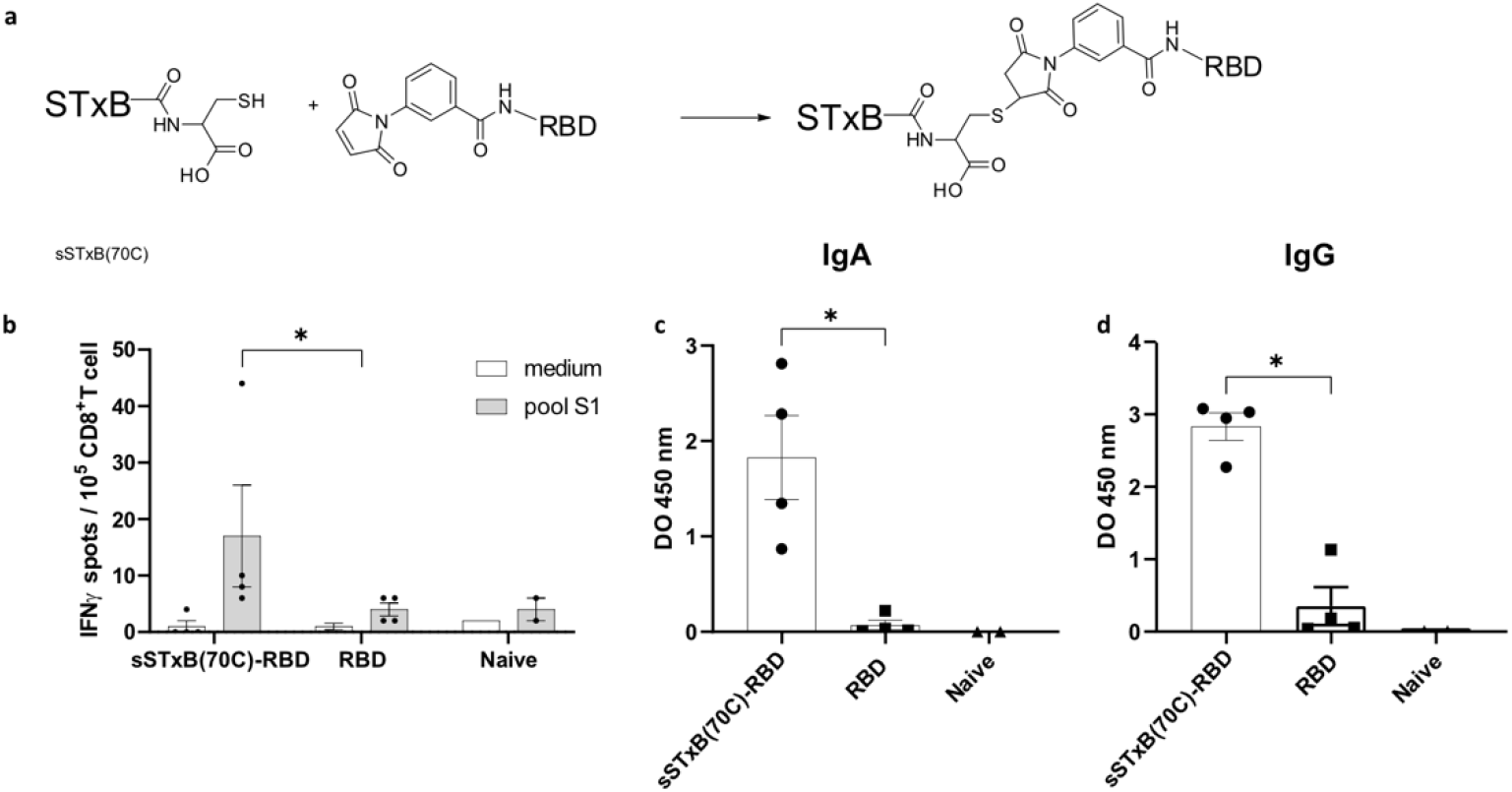
sSTxB(70C)-RBD induced higher RBD-specific CD8^+^ T cell and mucosal IgA responses in the lung than non-vectorized RBD. a) Conjugation of RBD to sSTxB(70C) through thiol-maleimide reaction. b-d) Balb/c mice (3-4 per group) were immunized at D0 and D14 via the intranasal route with 0.5 nmol of sSTxB(70C)-RBD or RBD, both combined with c-di-GMP as an adjuvant. Mice were sacrificed on day 21. Naive non-immunized mice were used as controls. After perfusion, the lungs were collected, and cells dissociated. CD8^+^ T cells were then purified and submitted to an IFNγ Elispot assay in which cells were sensitized or not (medium) with the S1 peptide pool encompassing the RBD sequence of SARS-CoV-2 (b). Bronchoalveolar lavage was also collected at day 21, diluted to ½, upon which specific anti-RBD IgA (c) and IgG (d) were measured. One out of 3 representative independent experiments is shown. Mean±SEM Mann-Whitney t-test. *P < 0.05.

**Figure 5:**
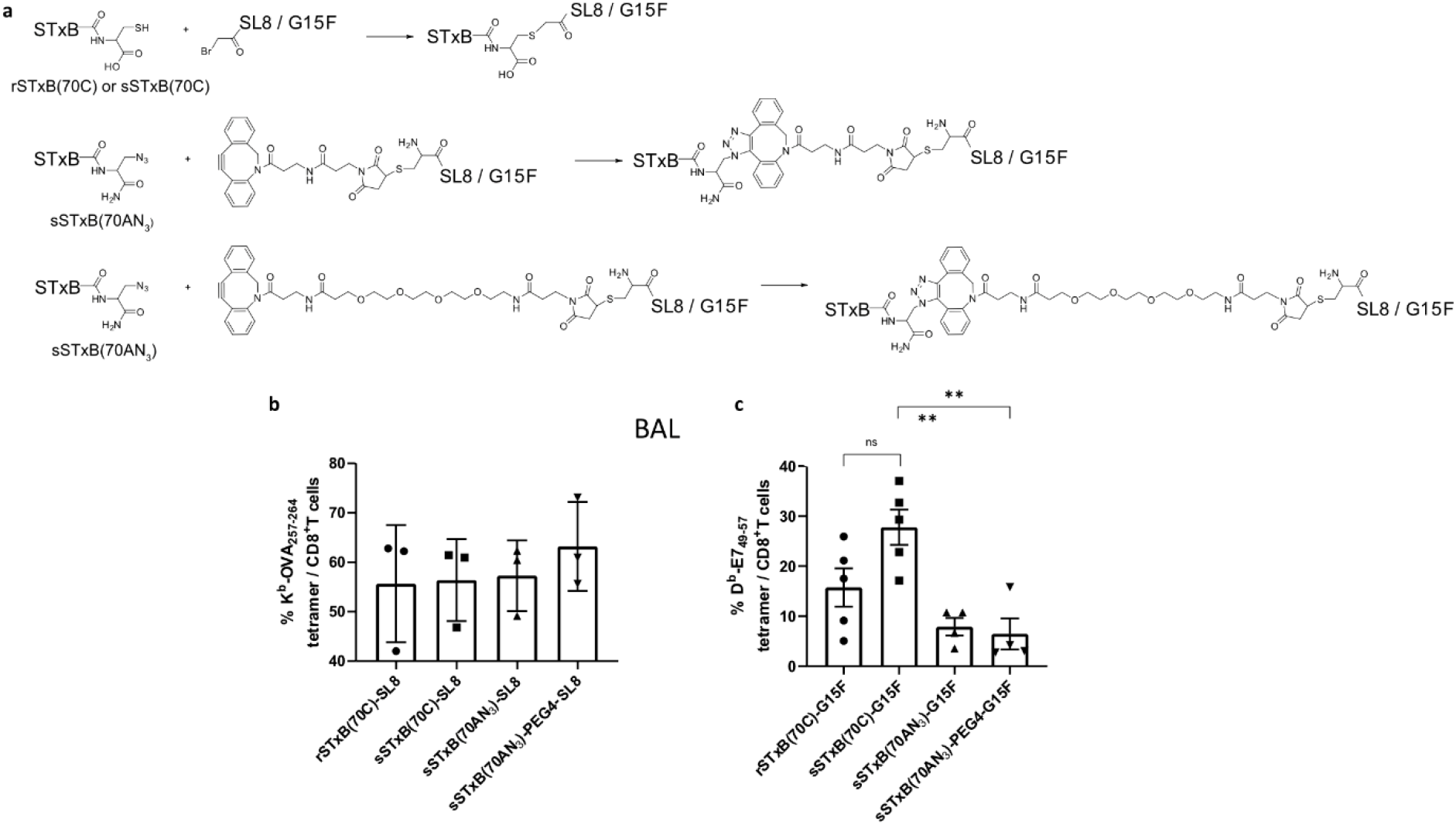
Various synthetic STxB conjugates induced mucosal-specific CD8^+^ T cells after vaccination in the C57Bl/6J mice model. a) Conjugation between rSTxB(70C), sSTxB(70C), or sSTxB(70AN_3_) and the antigenic peptides SL8 and G15F. b-c) Mice were immunized via the intranasal route at D0 and D14 with the indicated conjugates combined with the adjuvant α-GalCer (2 μg). On day 21, mice were sacrificed, and the broncho-alveolar lavage (BAL) was collected. Cells were labelled with anti-CD8 and specific K^b^-OVA_257-264_ (b) or D^b^-E7_49-57_ (c) tetramers. Results from 3-5 mice per group of one out of two representative experiments are shown. Background obtained from isotype control (D^b^-non-relevant peptide) was deduced (in general less than 1%). Mean±SEM Mann-Whitney t-test. *P < 0.05, ** P < 0.01.

For both antigenic peptides (SL8 and G15F), the induction of specific CD8^+^ T cells, as detected by flow cytometry analyses using specific tetramers, was as efficient or even better with synthetic STxB than with recombinant STxB (Figure 5). Between sSTxB variants, the performance levels were similar, except the sSTxB(70C)-G15F conjugate that stood out in its capacity of inducing high levels of anti-E7_49-57_ CD8^+^ T cells. Of note, CD8^+^ T cells that were induced in the lung had a memory resident phenotype (TRM), as characterized by CD103 expression alone or in combination with CD49a (Figure 6).

**Figure 6:**
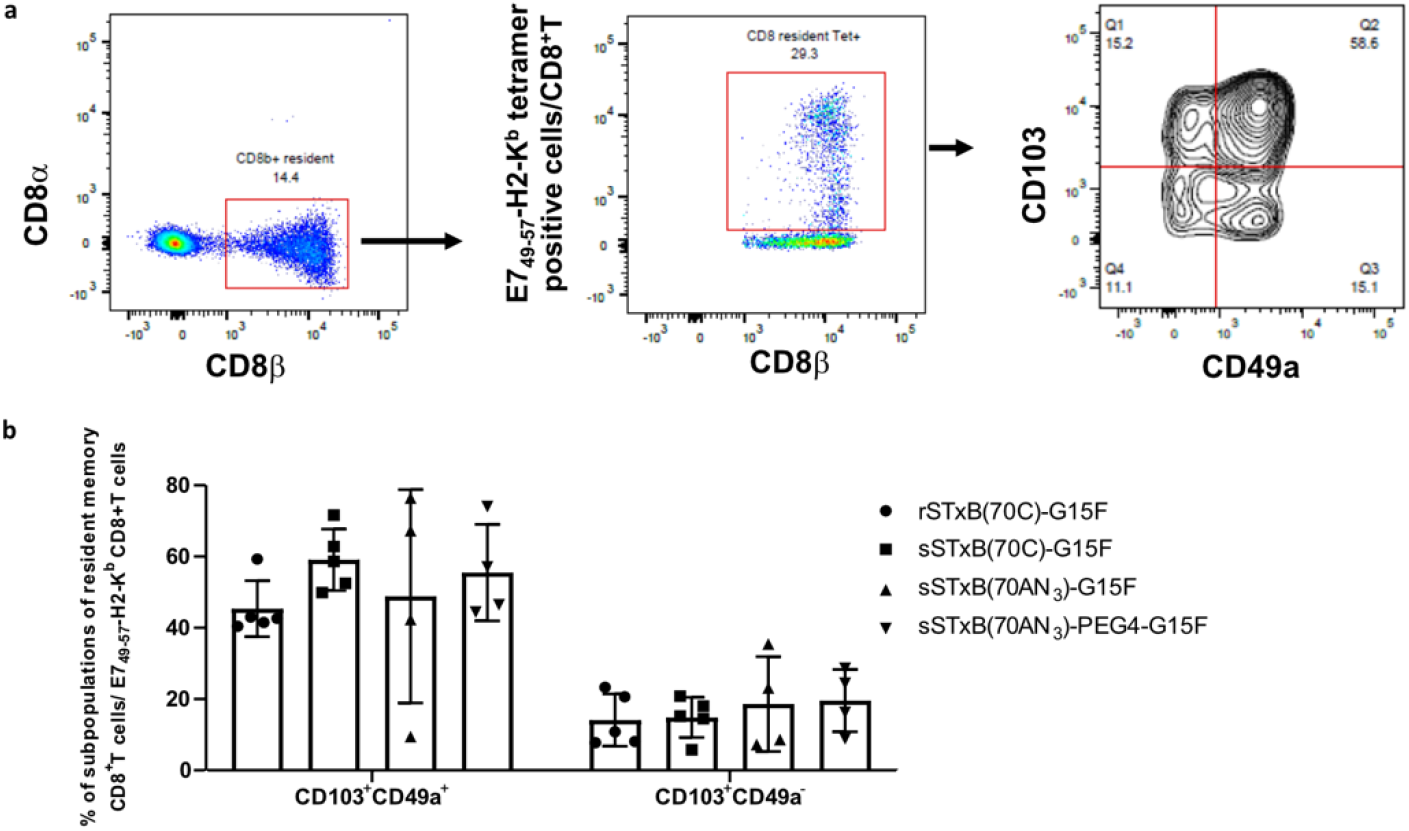
sSTxB(70C)-G15F induced resident memory CD8^+^ T cells in the lung. C57Bl/6J mice were immunized via the intranasal route at D0 and D14 with synthetic or recombinant STxB(70C)-G15F conjugates combined with the adjuvant α-GalCer (2 μg). At day 21, mice were sacrificed and the broncho-alveolar lavage (BAL) was collected. Cells were labeled with anti-CD8α, anti-CD8β, anti-CD103, anti-CD49a and D^b^-E7_49-57_ tetramers. a) Cells were first gated on CD8β^+^CD8α negative to eliminate NK and dendritic cells expressing CD8α. Then the CD8β^+^ tetramer^+^ T cells were analyzed for the expression of CD103 and CD49a which are biomarkers used to define resident memory T cells (TRM). Results from one out of five representative mice are shown. b) C57Bl/6J mice (n = 5) were immunized as described in Figure 5c with rSTxB(70C)-G15F or the indicated synthetic STxB conjugates with G15F. Means of subpopulations of specific resident memory CD8^+^ T cells defined by the expression of CD103 and/or CD49a are shown.

## 4. Conclusion

In this study, we have developed conditions for the linear synthesis of STxB without LC purification steps. STxB variants were synthesized and shown to be performant in inducing mucosal immunity in mice. The robustness of synthetic STxB as a mucosal vaccination vector was highlighted by its efficiency when coupled to different types of antigens, i.e., peptides (SL8, G15F) or full-size proteins (RBD), in different genetic backgrounds (Balb/c or C57bl/6J mice), and associated with different adjuvants (c-di-GMP, αGalCer). FDA legislation has recently evolved concerning synthetic peptides longer than 40 amino acids that are now classified as biologicals, which facilitates regulatory issues for projects that aim at developing synthetic proteins like STxB towards the clinics. The possibility of introducing several handles into STxB opens new avenues for the engineering of this protein, e.g., by site-specific modification by hydrophobic patches to further foster membrane translocation [36] and/or by concomitant coupling to antigens and adjuvants on the same STxB molecules for optimal cross-priming. Our study identifies a novel synthetic vaccine scaffold with multiple biomedical applications and development opportunities. The induction of functional CD8^+^ T cells with resident memory phenotype and mucosal IgA supports the development of STxB to address the still unmet medical need for a fully synthetic mucosal vaccine platform.

## Credit author statement

Anne Billet: Conceptualization; Data curation; Formal analysis; Writing - review & editing. Justine Hadjerci: Data curation; Formal analysis; Writing - original draft; Writing - review & editing. Thi Tran: Data curation; Formal analysis; Writing - review & editing. Pascal Kessler: Conceptualization; Data curation; Formal analysis; Writing - review & editing. Jonathan Ulmer: Data curation; Formal analysis. Gilles Mourier: Conceptualization; Data curation; Formal analysis. Marine Ghazarian: Data curation; Formal analysis. Anthony Gonzalez: Data curation; Formal analysis. Robert Thai: Data curation; Formal analysis. Pauline Urquia: Data curation; Formal analysis. Anne-Cécile Van Baelen: Data curation; Formal analysis. Annalisa Meola: Data curation; Formal analysis. Ignacio Fernandez: Data curation; Formal analysis. Stéphanie Deville-Foillard: Provided technical guidance on peptide synthesis; Writing - review & editing. Ewan MacDonald: Formal analysis. Léa Paolini: Data curation; Formal analysis. Frédéric Schmidt: Conceptualization. Félix A. Rey: Project administration; Resources; Supervision; Writing - review & editing. Michael S. Kay: Provided technical guidance on STxB synthesis. Eric Tartour: Conceptualization; Funding acquisition; Investigation; Project administration; Supervision; Writing - review & editing. Denis Servent: Conceptualization; funding acquisition; Investigation; Project administration; Supervision; Writing - review & editing. Ludger Johannes: Conceptualization; Funding acquisition; Investigation; Project administration; Supervision; Writing - original draft; Writing - review & editing.

## Acknowledgments

This study was supported by grants from Mizutani Foundation (reference n° 200014 to L.J.), Institut National du Cancer (INCa contract n°2019-1-PLBIO-05-1 to L.J., E.T. D.S., and INCa Contract PLBIO22-147 to E.T.), La Ligue Nationale contre le Cancer (convention N° AAPARN 2021.LCC/ChP to L.J. and E.T.), Equipes Labellisées Fondation pour la Recherche Médicale (EQU202103012926 to L.J.) and Ligue contre le Cancer (EL 2020 to E.T.). F.A.R. received funding from the Institut Pasteur Coronavirus Task Force (Allospike project), the CNRS and grant ANR-10-LABX-62-IBEID. A.B. was funded by the French Ministry of Higher Education and Research (AMX funding) and La Ligue Nationale Contre le Cancer, and received support from “Frontières de l’Innovation en Recherche et Éducation” (FIRE) Doctoral School – Bettencourt Program. J.H. was funded by the Fondation pour la Recherche Médicale. This work was also funded by the Agence Nationale de la Recherche (Labex Immuno-Oncology to E.T.) and the Institut National du Cancer (Grant SIRIC CARPEM to E.T.). This work was supported by a Curie Innov grant from Institut Carnot Curie Cancer (ANR 18 CARN 0009 01) to LJ. We acknowledge Raphaël Rodriguez’s team from Institut Curie for the use of various instruments and the Cell and Tissue Imaging (PICT-IBiSA) and Nikon Imaging Centre, Institut Curie, member of the French National Research Infrastructure France-BioImaging (ANR10-INBS-04). We greatly acknowledge Patrick England and Sébastien Brûlé from the Biophysics Platform at Institut Pasteur for their help in the biophysical characterization of STxB.

## Supplementary data

Supplementary data to this article can be found online.

## Data availability

Data will be made available on request.

