## Supplementary Data for "A synthetic delivery vector for mucosal vaccination"

### Supplementary Tables and Figures

### Linear full-length STxB synthesis

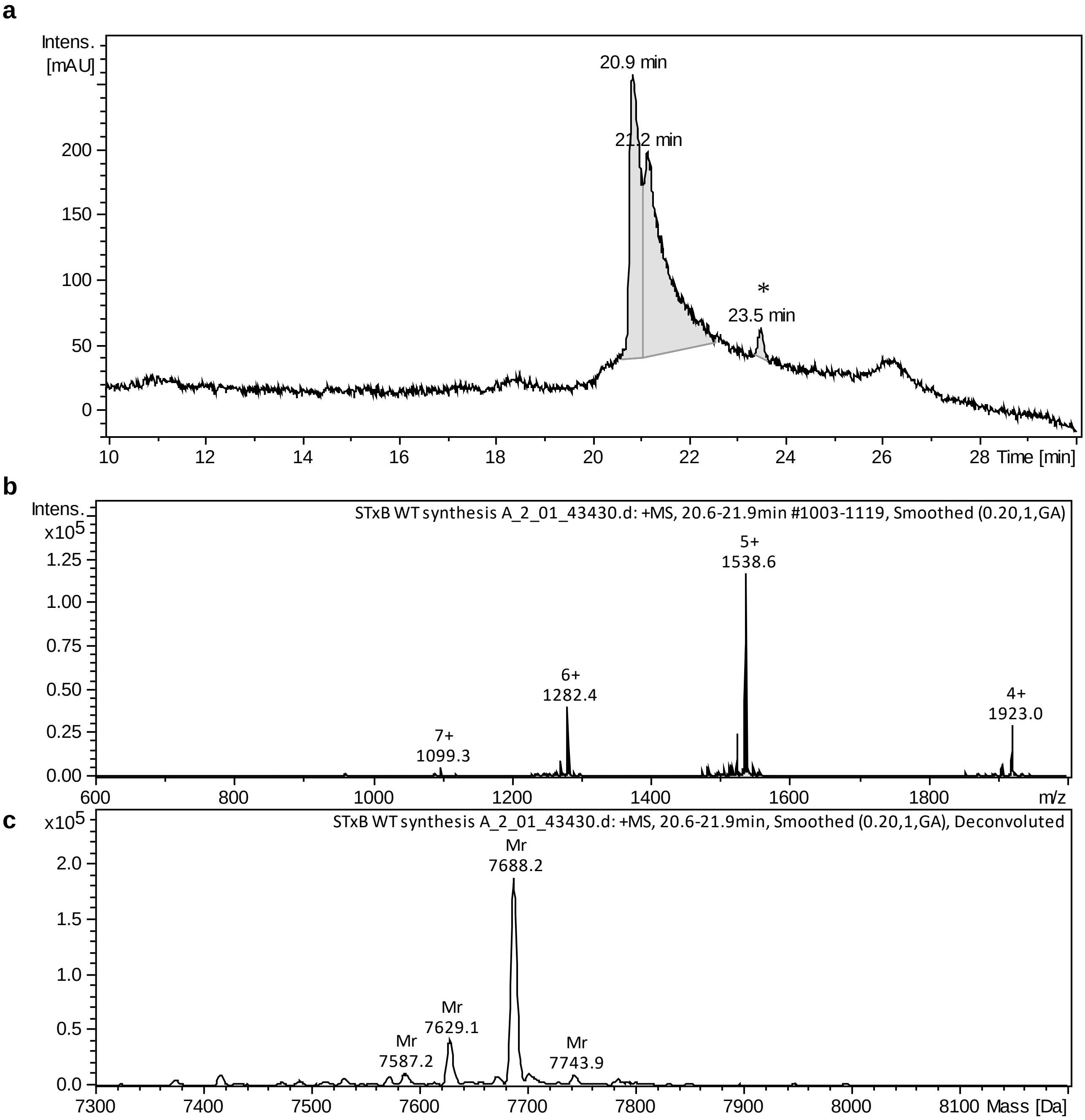

**Figure S1:** HPLC-MS of folded STxB synthesised following condition A (see Table 1).
a) HPLC chromatogram (λ = 214 nm). b) Mass spectrum (ESI+). c) Deconvoluted averaged MS spectra. (*): Traces of undefined contaminant.

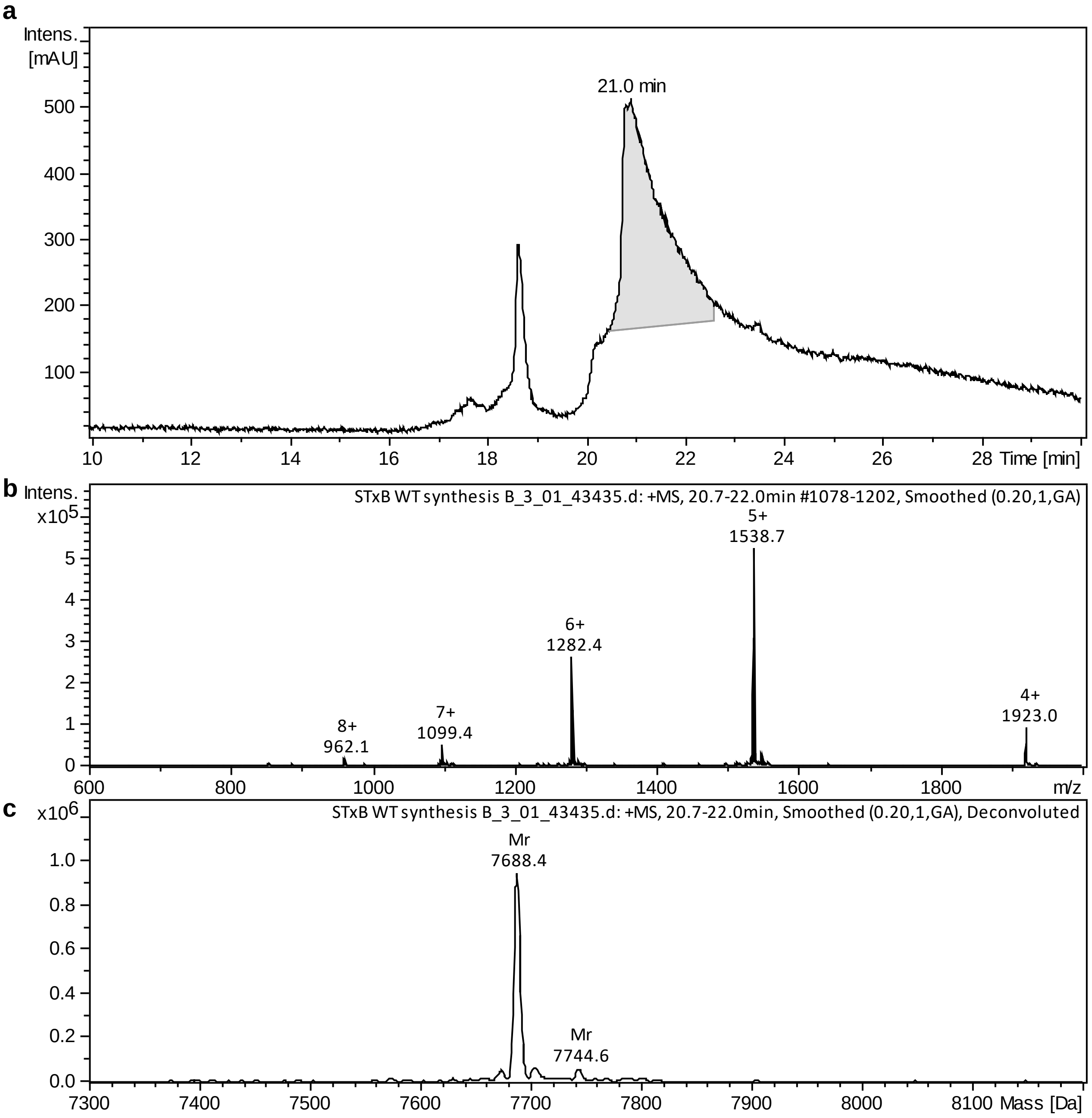

**Figure S2:** HPLC-MS of folded STxB synthesised following condition B (see Table 1).
a) HPLC chromatogram (λ = 214 nm). b) Mass spectrum (ESI+). c) Deconvoluted averaged MS spectra.

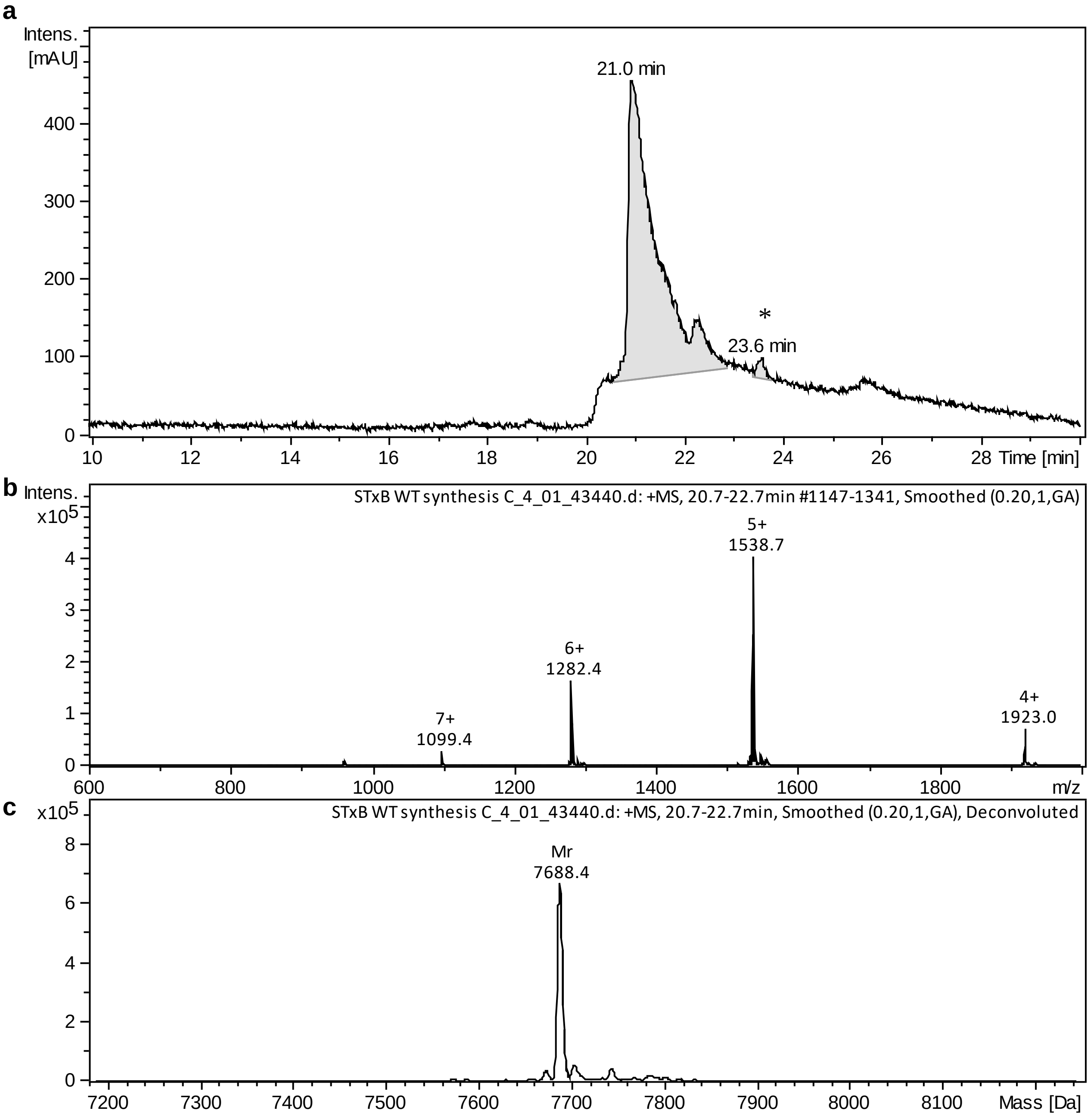

**Figure S3:** HPLC-MS of folded STxB synthesised following condition C (see Table 1).

a) HPLC chromatogram (λ = 214 nm). b) Mass spectrum (ESI+). c) Deconvoluted averaged MS spectra. (*): Traces of undefined contaminant.

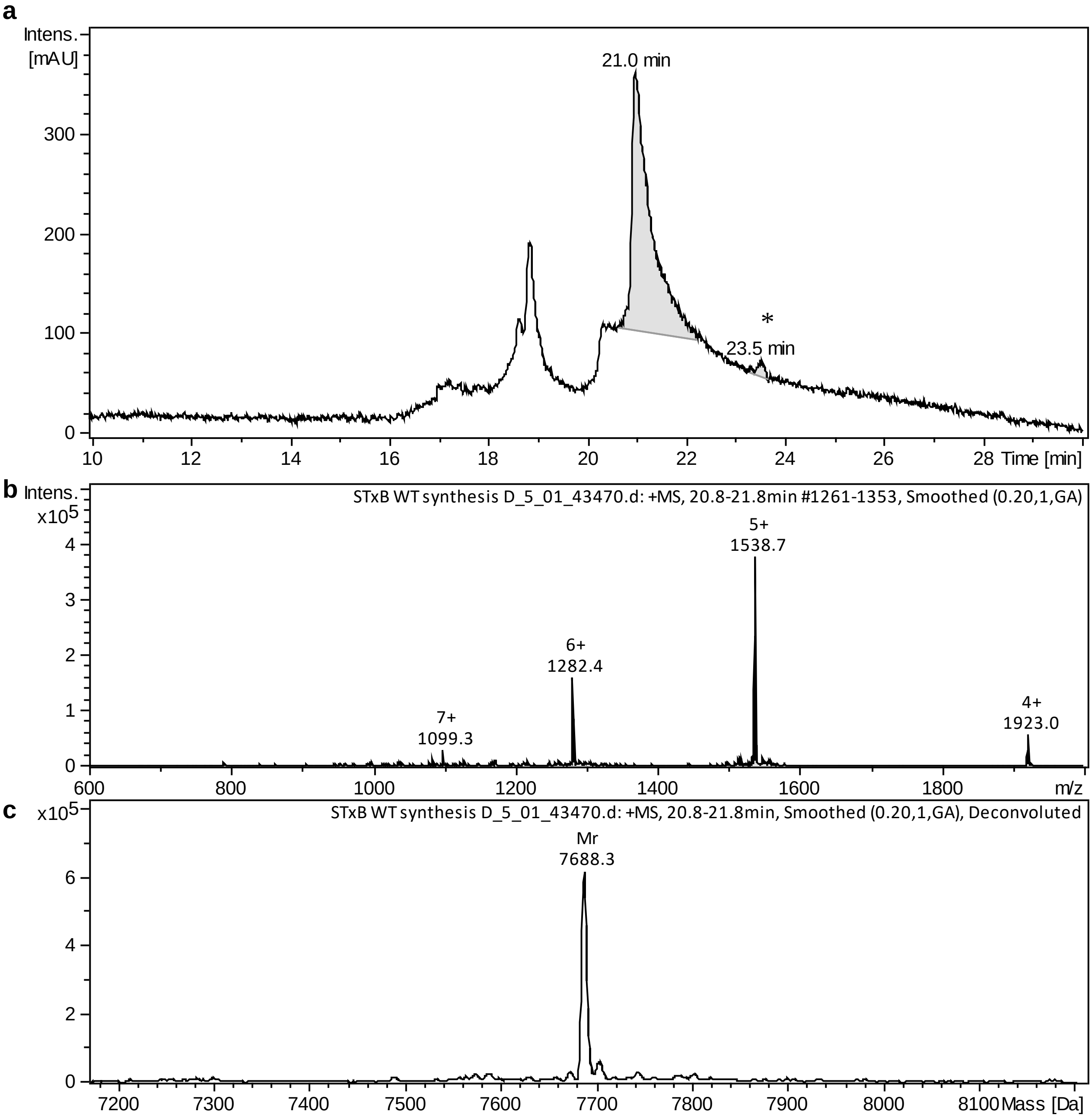

**Figure S4:** HPLC-MS of folded STxB synthesised following condition D (see Table 1).

a) HPLC chromatogram (λ = 214 nm). b) Mass spectrum (ESI+). c) Deconvoluted averaged MS spectra. (*): Traces of undefined contaminant.

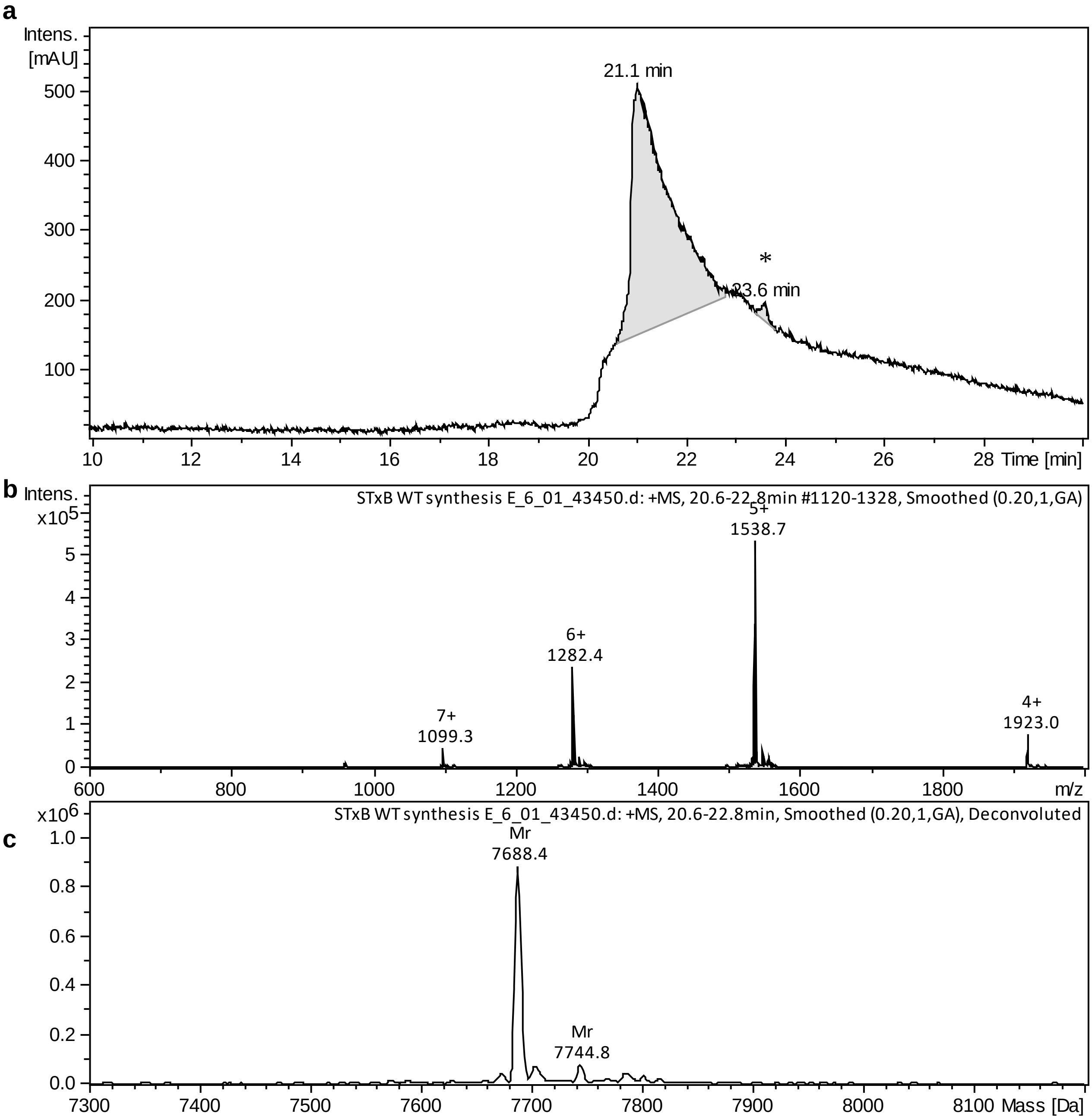

**Figure S5:** HPLC-MS of folded STxB synthesised following condition E (see Table 1).

a) HPLC chromatogram (λ = 214 nm). b) Mass spectrum (ESI+). c) Deconvoluted averaged MS spectra. (*): Traces of undefined contaminant.

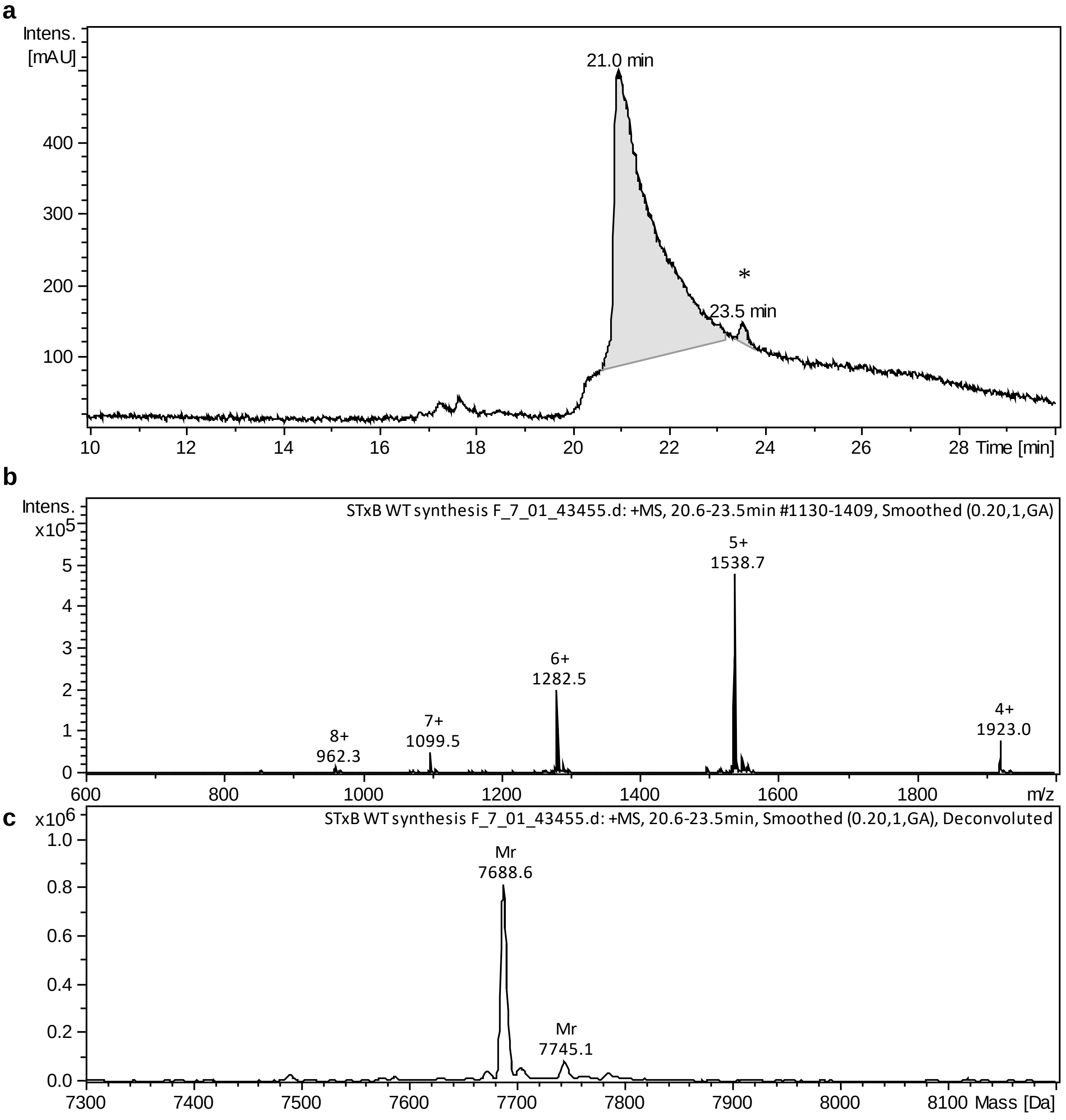

**Figure S6:** HPLC-MS of folded STxB synthesised following condition F (see Table 1).

a) HPLC chromatogram (λ = 214 nm). b) Mass spectrum (ESI+). c) Deconvoluted averaged MS spectra. (*): Traces of undefined contaminant.

a)

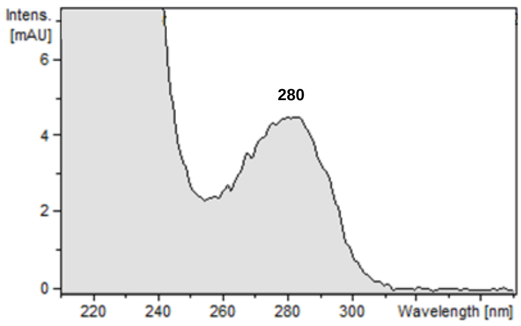

b)

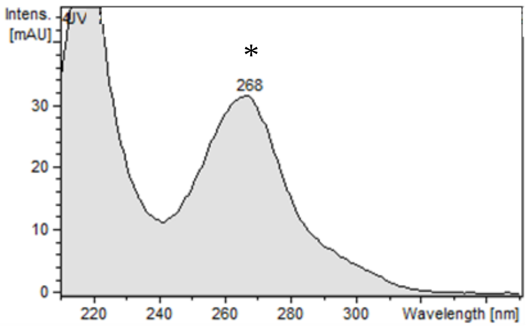

**Figure S7:** (*) Traces of undefined contaminant at 23.5 min.
a) UV spectrum corresponding to folded STxB ($\lambda_{max}$ = 280 nm). b) UV spectrum of the peak at 23.5 min corresponding to an undefined contaminant. The UV spectrum of the peak at 23.5 min seems to not correspond to a protein compound ($\lambda_{max}$ = 280 nm) with an unexpected $\lambda_{max}$ at 268 nm.

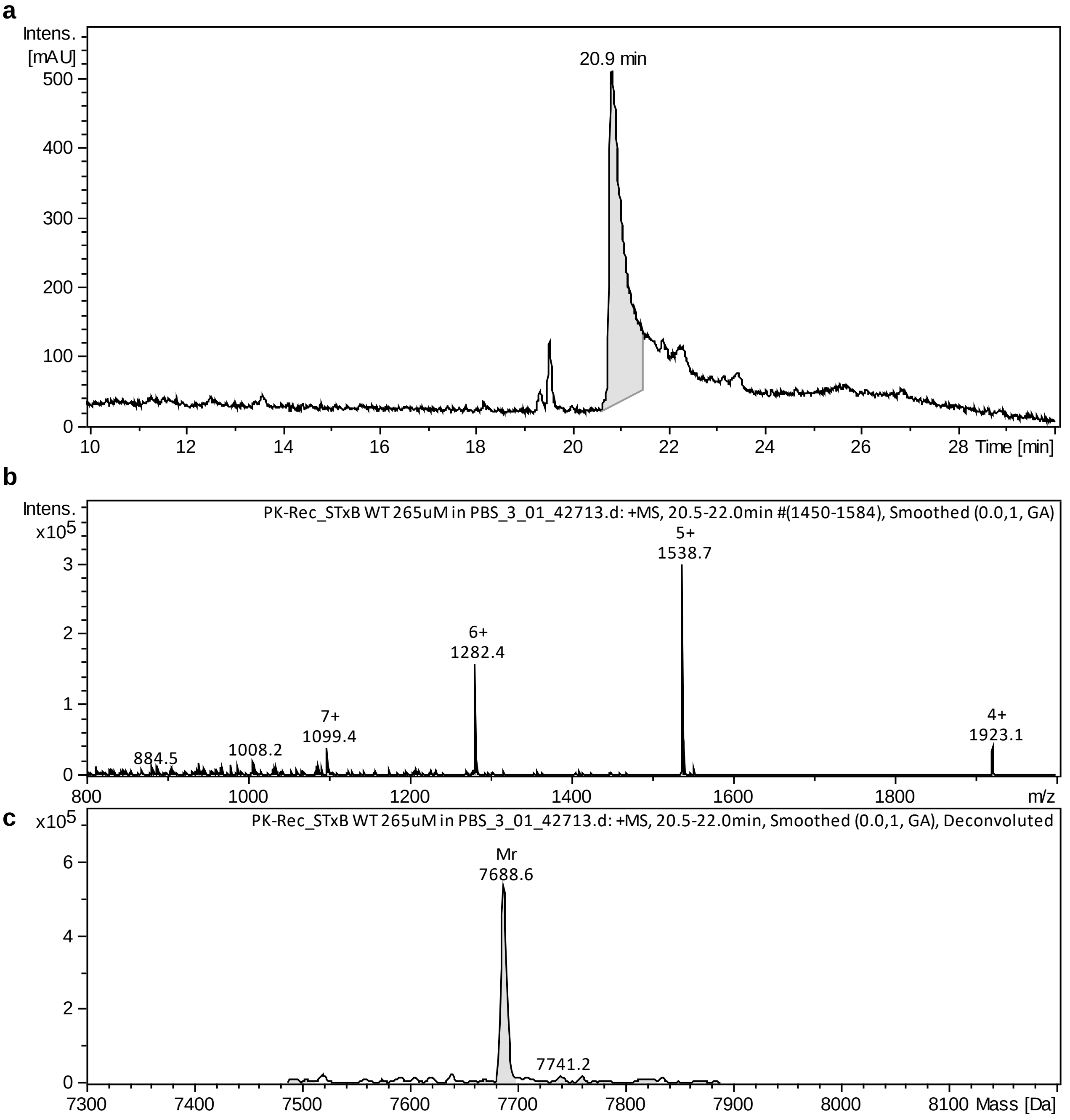

**Figure S8:** HPLC-MS of recombinant STxB.

a) HPLC chromatogram (λ = 214 nm). b) Mass spectrum (ESI+). c) Deconvoluted averaged MS spectra.

MS m/z $C_{339}H_{528}N_{90}O_{108}S_3$ $[M+H^+]^+$ calculated: 7688.6, found: 7688.6; $[M+4H^+]^{4+}$ calculated: 1923.1, found: 1923.1; $[M+5H^+]^{5+}$ calculated: 1538.7, found: 1538.7; $[M+6H^+]^{6+}$ calculated: 1282.4, found: 1282.4, $[M+7H^+]^{7+}$ calculated: 1099.4, found: 1099.4.

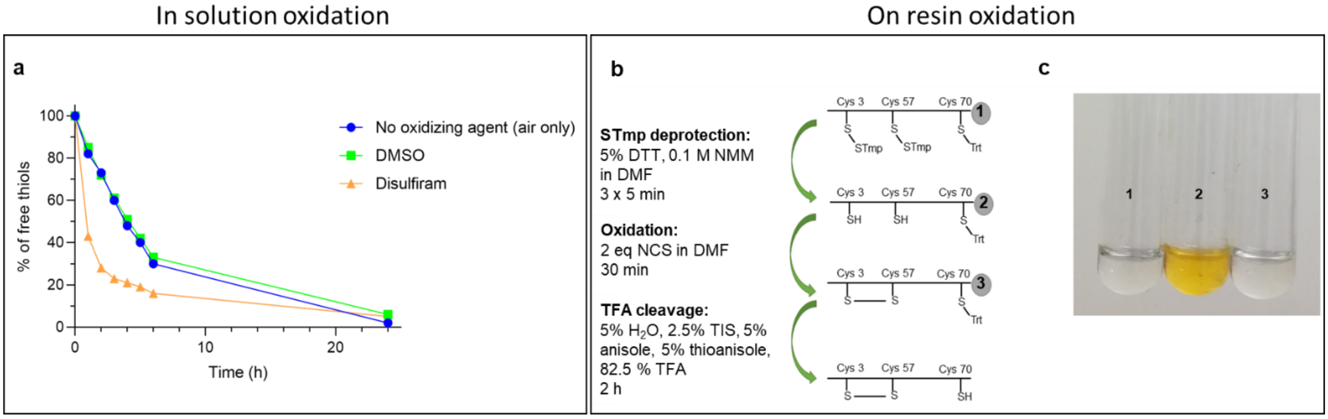

**Figure S9:** Oxidation optimization.

a) In solution oxidation kinetics of crude synthetic STxB with or without the addition of oxidizing agents (DMSO: 2 %; disulfiram: 2 eq) under stirring at 37 °C (followed using Ellman's test in 7 M GndHCl, 50 mM phosphate pH 8). Percentages are expressed in comparison to free thiols at t = 0 h. b-c) On resin oxidation of sSTxB(70C): free thiol detection on a few beads withdrawn at different steps. The detection was performed using 2,2'-dithiobis-(5-nitropyridine) (DTNP), which liberated a yellow product after reaction with free thiols. 1: Beads after SPPS; 2: Beads after STmp deprotection; 3: Beads after oxidation.

Abbreviations: Trt: trityl, STmp: trimethoxyphenylthio, DTT: dithiothreitol, NMM: *N*-methylmorpholine, DMF: dimethylformamide, NCS: *N*-chlorosuccinimide, TIS: triisopropylsilane, TFA: trifluoroacetic acid.

### Chemical synthesis opens new possibilities for bioorthogonal conjugations with STxB

**Table S1:** Synthesis yields and functionality of STxB azido variants.
When not specified otherwise, the variants had a C-terminal -COOH.

| Azido variant | Azide localization on STxB (in red). Gb3 analogs (in black) | Fmoc synthesis yield (%) | Crude peptide mass (mg) | Oxidation and folding yield (%) | Overall Yield (%) | Retrograde trafficking assay | Conjugation with DBCO |
|---|---|---|---|---|---|---|---|
| STxB(T1A$N_3$) | 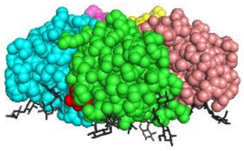 | 69 | 54.5 | 35 | 24 | Less signal | Validated |
| STxB(D3K$N_3$) | 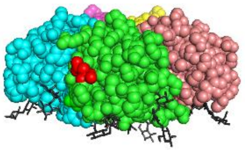 | 69 | 43.8 | 31 | 21 | Functional | Validated |
| STxB(T6K$N_3$) | 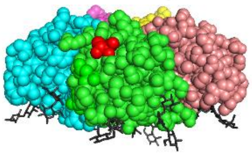 | 42 | 63.7 | 16 | 7 | Functional | Validated |
| STxB(K8K$N_3$) | 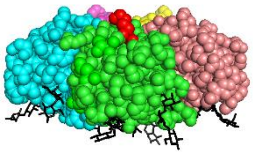 | 56 | 77.3 | 36 | 20 | Functional | Validated |

| Azido variant | Azide localization on STxB (in red). Gb3 analogs (in black) | Fmoc synthesis yield (%) | Crude peptide mass (mg) | Oxidation and folding yield (%) | Overall Yield (%) | Retrograde trafficking assay | Conjugation with DBCO |
|---|---|---|---|---|---|---|---|
| STxB(E10K$N_3$) | 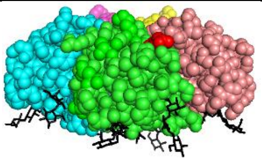 | 70 | 72.9 | 30 | 21 | Functional | Validated |
| STxB(Y11K$N_3$) | 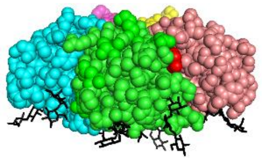 | 58 | 73.1 | 29 | 17 | Functional | Validated |
| STxB(Y11Phe(4-$CH_2$-$N_3$)) | 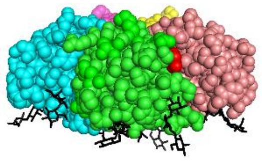 | 43 | 63.5 | 28 | 12 | Functional | Validated |
| STxB(K23K$N_3$) | 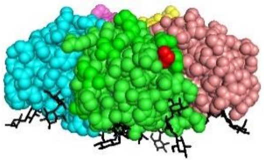 | / | 55 | 38 | / | Functional | Validated |
| STxB(D26K$N_3$) | 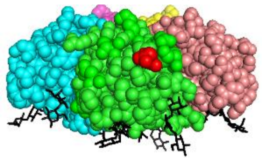 | 59 | 81.1 | 12 | 7 | Functional | Product formed, but low concentration |
| STxB(K27K$N_3$) | 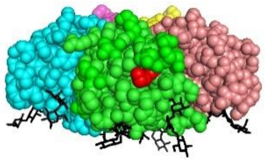 | 57 | 64.4 | 33 | 19 | Functional | Validated |
| STxB(T49K$N_3$) | 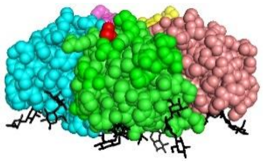 | 51 | 92.5 | 30 | 15 | Functional | Validated |
| STxB(K53K$N_3$) | 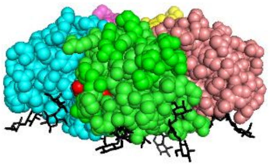 | 50 | 71.7 | 26 | 13 | Functional | Validated |
| STxB(H58A$N_3$) | 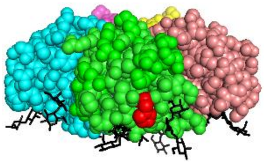 | 44 | 64 | 29 | 13 | Functional [1] | Validated |
| STxB(N59K$N_3$) | 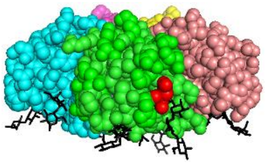 | 45 | 68.5 | 27 | 12 | Functional | Validated |

| Azido variant | Azide localization on STxB (in red). Gb3 analogs (in black) | Fmoc synthesis yield (%) | Crude peptide mass (mg) | Oxidation and folding yield (%) | Overall Yield (%) | Retrograde trafficking assay | Conjugation with DBCO |
|---|---|---|---|---|---|---|---|
| STxB(R69K$N_3$)-$CONH_2$ | 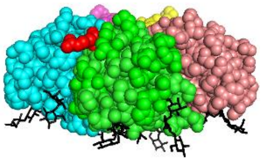 | 45 | 61.1 | 22 | 10 | Functional | Validated |
| STxB(70A$N_3$)-$CONH_2$ | 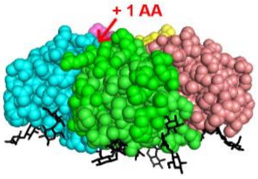  | / | 18.6 | 29 | / | Functional | Validated |
| STxB(70K$N_3$)-$CONH_2$ | 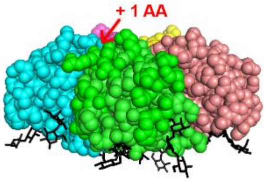  | 61 | 25.3 | 21 | 13 | Functional | Validated |

[1] After direct labelling with Cy3, not recognized by 13C4 Ab.

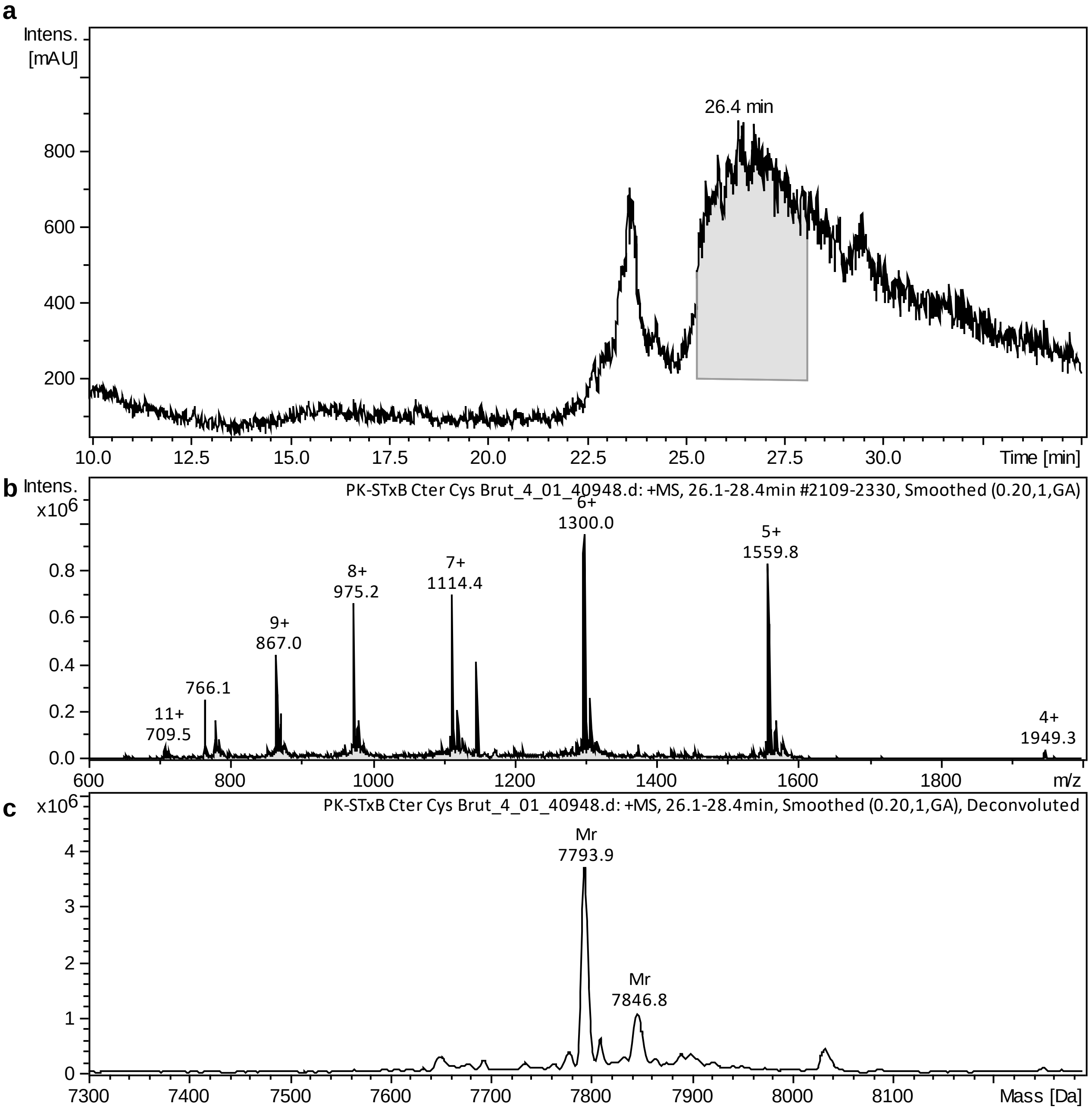

**Figure S10:** HPLC-MS of crude sSTxB(70C) peptide.

a) HPLC chromatogram (λ = 214 nm). b) Mass spectrum (ESI+). c) Deconvoluted averaged MS spectra. MS m/z $C_{342}H_{533}N_{91}O_{109}S_4$ Average MW calculated: 7793.8, found: 7793.9; $[M+4H^+]^{4+}$ calculated: 1949.5, found: 1949.3; $[M+5H^+]^{5+}$ calculated: 1559.8, found: 1559.8; $[M+6H^+]^{6+}$ calculated: 1300.0, found: 1300.0; $[M+7H^+]^{7+}$ calculated: 1114.4, found: 1114.4; $[M+8H^+]^{8+}$ calculated: 975.2, found: 975.2; $[M+9H^+]^{9+}$ calculated: 867.0, found: 867.0; $[M+11H^+]^{11+}$ calculated: 709.5, found: 709.5.

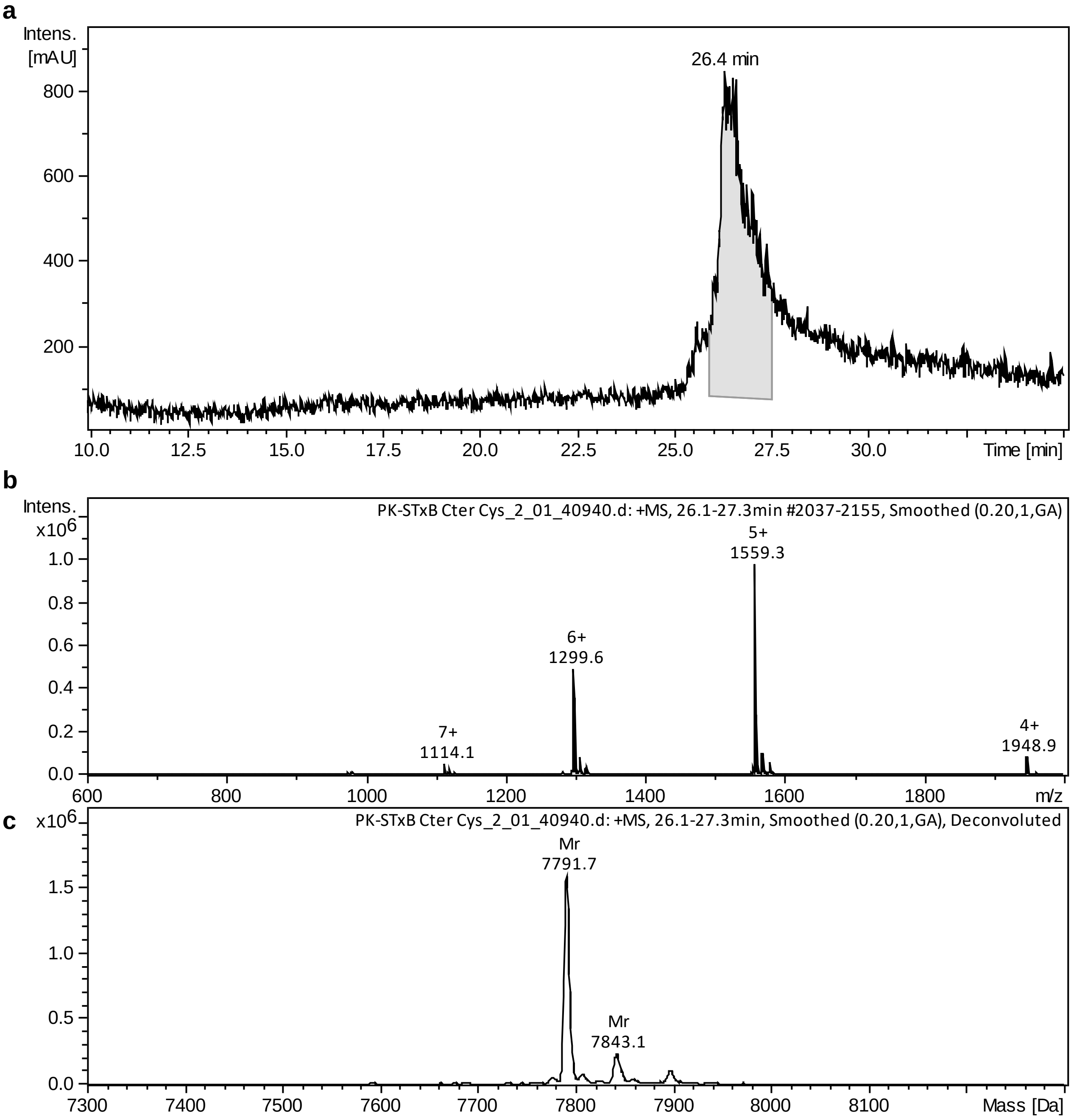

**Figure S11:** HPLC-MS of folded sSTxB(70C) protein.

a) HPLC chromatogram (λ = 214 nm). b) Mass spectrum (ESI+). c) Deconvoluted averaged MS spectra. MS m/z $C_{342}H_{533}N_{91}O_{109}S_4$ Average MW calculated: 7791.9, found: 7791.7; $[M+4H^+]^{4+}$ calculated: 1949.0, found: 1948.9; $[M+5H^+]^{5+}$ calculated: 1559.4, found: 1559.3; $[M+6H^+]^{6+}$ calculated: 1299.7, found: 1299.6; $[M+7H^+]^{7+}$ calculated: 1114.1, found: 1114.1.

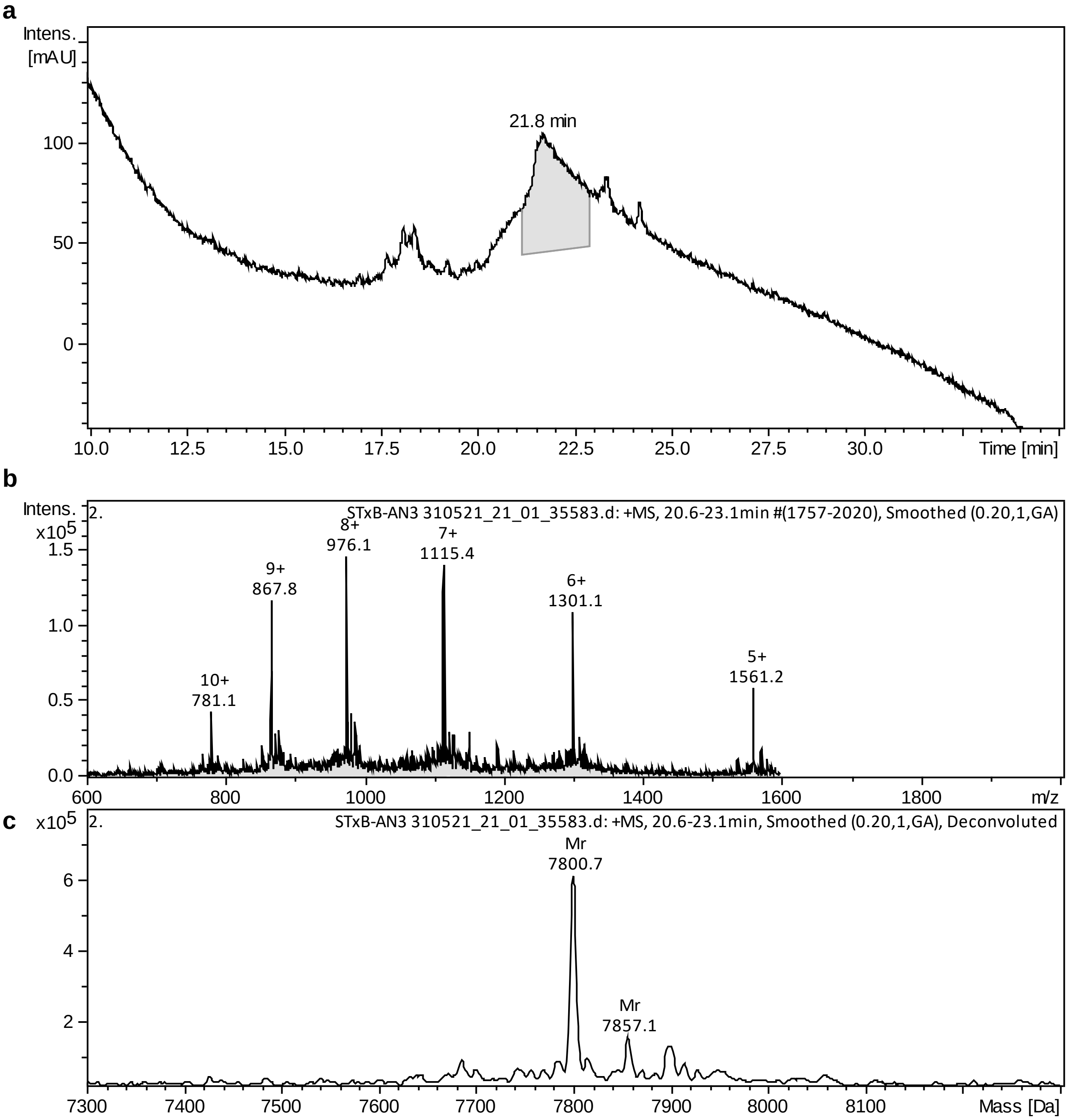

**Figure S12:** HPLC-MS of crude sSTxB(70AN$_3$) peptide.

a) HPLC chromatogram (λ = 214 nm). b) Mass spectrum (ESI+). c) Deconvoluted averaged MS spectra. MS m/z $C_{342}H_{533}N_{94}O_{109}S_3$ Average MW calculated: 7800.7, found: 7800.7; $[M+5H^+]^{5+}$ calculated: 1561.2, found: 1561.2; $[M+6H^+]^{6+}$ calculated: 1301.2, found: 1301.1; $[M+7H^+]^{7+}$ calculated: 1115.4, found: 1115.4; $[M+8H^+]^{8+}$ calculated: 976.1, found: 976.1; $[M+9H^+]^{9+}$ calculated: 867.8, found: 867.8; $[M+10H^+]^{10+}$ calculated: 781.1, found: 781.1.

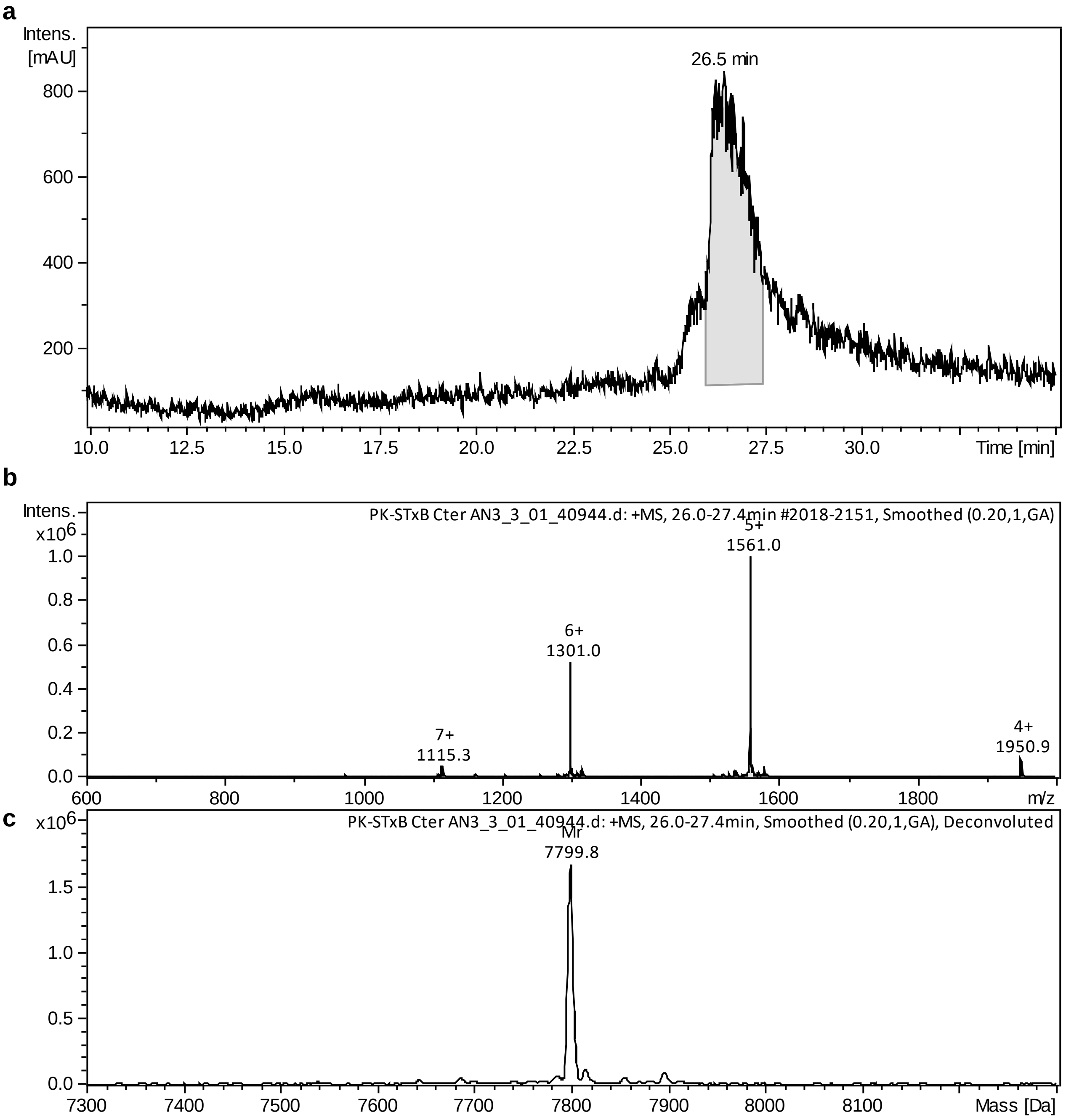

**Figure S13:** HPLC-MS of folded sSTxB(70AN$_3$) protein.

a) HPLC chromatogram (λ = 214 nm). b) Mass spectrum (ESI+). c) Deconvoluted averaged MS spectra. MS m/z $C_{342}H_{533}N_{94}O_{109}S_3$ Average MW calculated: 7798.7, found: 7799.8; $[M+4H^+]^{4+}$ calculated: 1950.7, found: 1950.9; $[M+5H^+]^{5+}$ calculated: 1560.7, found: 1561.0; $[M+6H^+]^{6+}$ calculated: 1300.8, found: 1301.0; $[M+7H^+]^{7+}$ calculated: 1115.1, found: 1115.3.

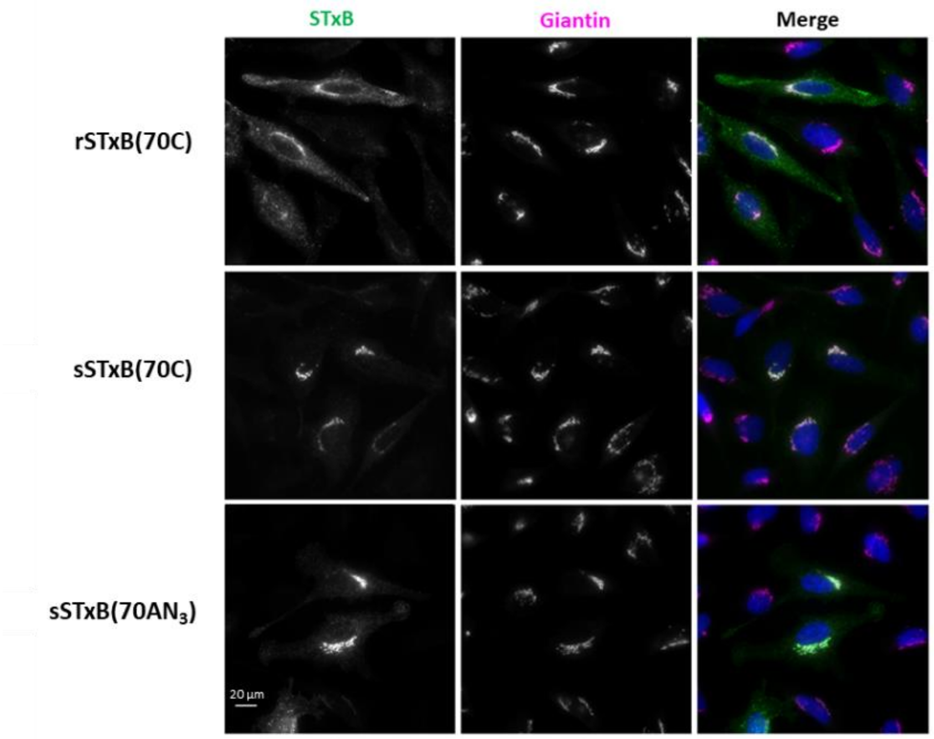

**Figure S14:** Retrograde trafficking of recombinant and synthetic STxB(70C) and sSTxB(70AN$_3$) on HeLa cells. For each variant, 40 nM of STxB was incubated for 30 min at 4 °C with cells, followed by PBS washes and incubation for 50 min at 37 °C for synchronized internalization. Merge: STxB in green, Golgi in magenta, and DNA dye Hoechst in blue. Scale bar: 20 µm.

### Synthetic STxB: an efficient delivery tool for mucosal vaccination

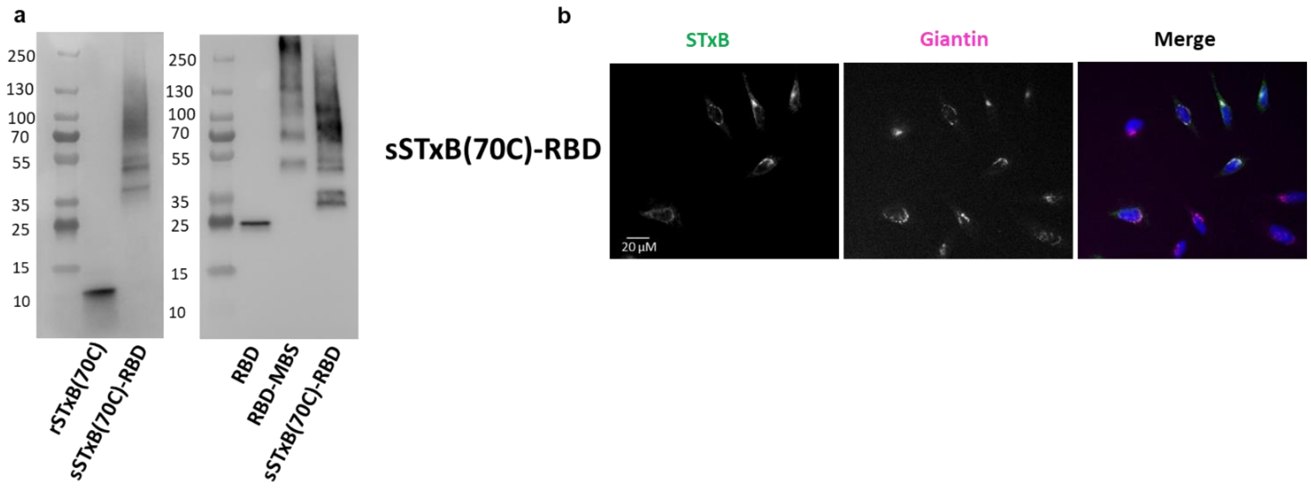

**Figure S15:** Characterization of sSTxB(70C)-RBD conjugate.

a) Western blot of sSTxB(70C)-RBD conjugate under denaturing conditions. Anti-STxB labeling (left) and anti-RBD labeling (right). Corresponding sizes for sSTxB(70C) monomer are: 7.8 kDa, and RBD: 22 kDa. Note that RBD is glycosylated. b) Retrograde trafficking of sSTxB(70C)-RBD conjugate on HeLa cells. 40 nM of sSTxB(70C)-RBD conjugate was incubated with cells for 30 min at 4 °C, followed by PBS washes and incubation for 50 min at 37 °C for synchronized internalization. Merge: STxB in green, Golgi in magenta, and DNA dye Hoechst in blue. Scale bar: 20 µm.

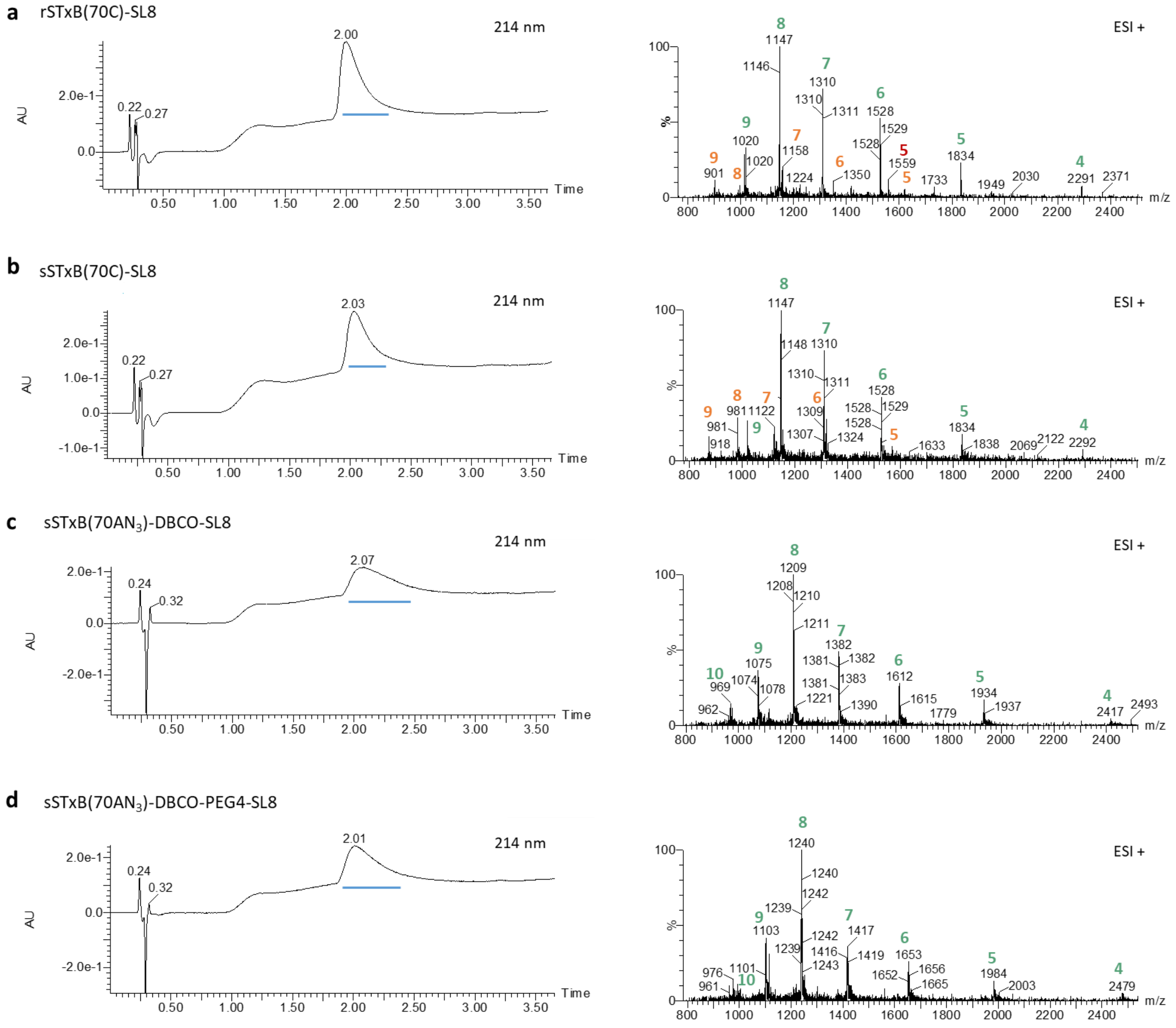

**Figure S16:** UPLC-MS analyses of sSTxB(70C) and sSTxB(70AN$_3$) conjugates with SL8.

Mass spectra (ESI+): Peaks corresponding to the expected product are annotated with their corresponding positive charge in green. Peaks annotated in orange correspond to monomers with a non-reactive cysteine (either due to a glutathione adduct on recombinant STxB, or to a *t*Bu adduct on synthetic STxB). Peaks annotated in red correspond to unconjugated reactive monomers.

a) $C_{405}H_{634}N_{106}O_{128}S_4$ $[M+4H^+]^{4+}$ calculated: 2292.5, found: 2291.5; $[M+5H^+]^{5+}$ calculated: 1834.2, found: 1833.7; $[M+6H^+]^{6+}$ calculated: 1528.6, found: 1528.2; $[M+7H^+]^{7+}$ calculated: 1310.4, found: 1310.2; $[M+8H+]^{8+}$ calculated: 1146.7, found: 1146.9; $[M+9H^+]^{9+}$ calculated: 1019.4, found: 1019.7.

b) $C_{405}H_{634}N_{106}O_{128}S_4$ $[M+4H^+]^{4+}$ calculated: 2292.5, found: 2292.0; $[M+5H^+]^{5+}$ calculated: 1834.2, found: 1834.0; $[M+6H^+]^{6+}$ calculated: 1528.6, found: 1528.5; $[M+7H^+]^{7+}$ calculated: 1310.4, found: 1310.1; $[M+8H^+]^{8+}$ calculated: 1146.7, found: 1146.7; $[M+9H^+]^{9+}$ calculated: 1019.4, found: 1019.8.

c) $C_{431}H_{659}N_{112}O_{133}S_4$ $[M+4H^+]^{4+}$ calculated: 2417.0, found: 2417.5; $[M+5H^+]^{5+}$ calculated: 1933.8, found: 1933.5; $[M+6H^+]^{6+}$ calculated: 1611.6, found: 1611.8; $[M+7H^+]^{7+}$ calculated: 1381.5, found: 1381.7; $[M+8H^+]^{8+}$ calculated: 1209.0, found: 1208.8; $[M+9H^+]^{9+}$ calculated: 1074.8, found: 1075.2; $[M+10H^+]^{10+}$ calculated: 967.4, found: 969.1.

d) $C_{442}H_{680}N_{113}O_{138}S_4$ $[M+4H^+]^{4+}$ calculated: 2478.8, found: 2479.0; $[M+5H^+]^{5+}$ calculated: 1983.2, found: 1983.5; $[M+6H^+]^{6+}$ calculated: 1652.8, found: 1652.7; $[M+7H^+]^{7+}$ calculated: 1416.9, found: 1416.9; $[M+8H^+]^{8+}$ calculated: 1239.9, found: 1240.0; $[M+9H^+]^{9+}$ calculated: 1102.2, found: 1102.9; $[M+10H^+]^{10+}$ calculated: 992.1, found: 993.4.

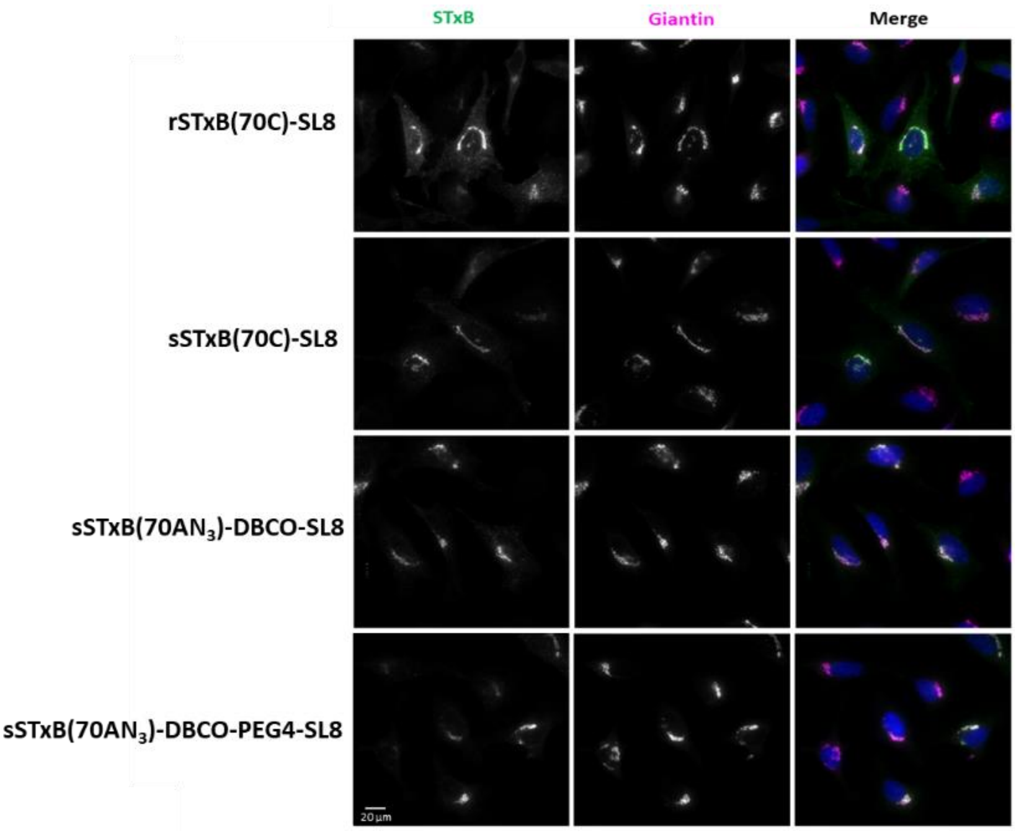

**Figure S17:** Retrograde trafficking on HeLa cells of sSTxB(70C) and sSTxB(70AN$_3$) conjugated with SL8. Cells were incubated with the indicated STxB conjugates for 30 min at 4 °C, followed by PBS washes and incubation for 50 min at 37 °C for synchronized internalization. Merge: STxB in green, Golgi in magenta, and DNA dye Hoechst in blue. Scale bar: 20 µm.

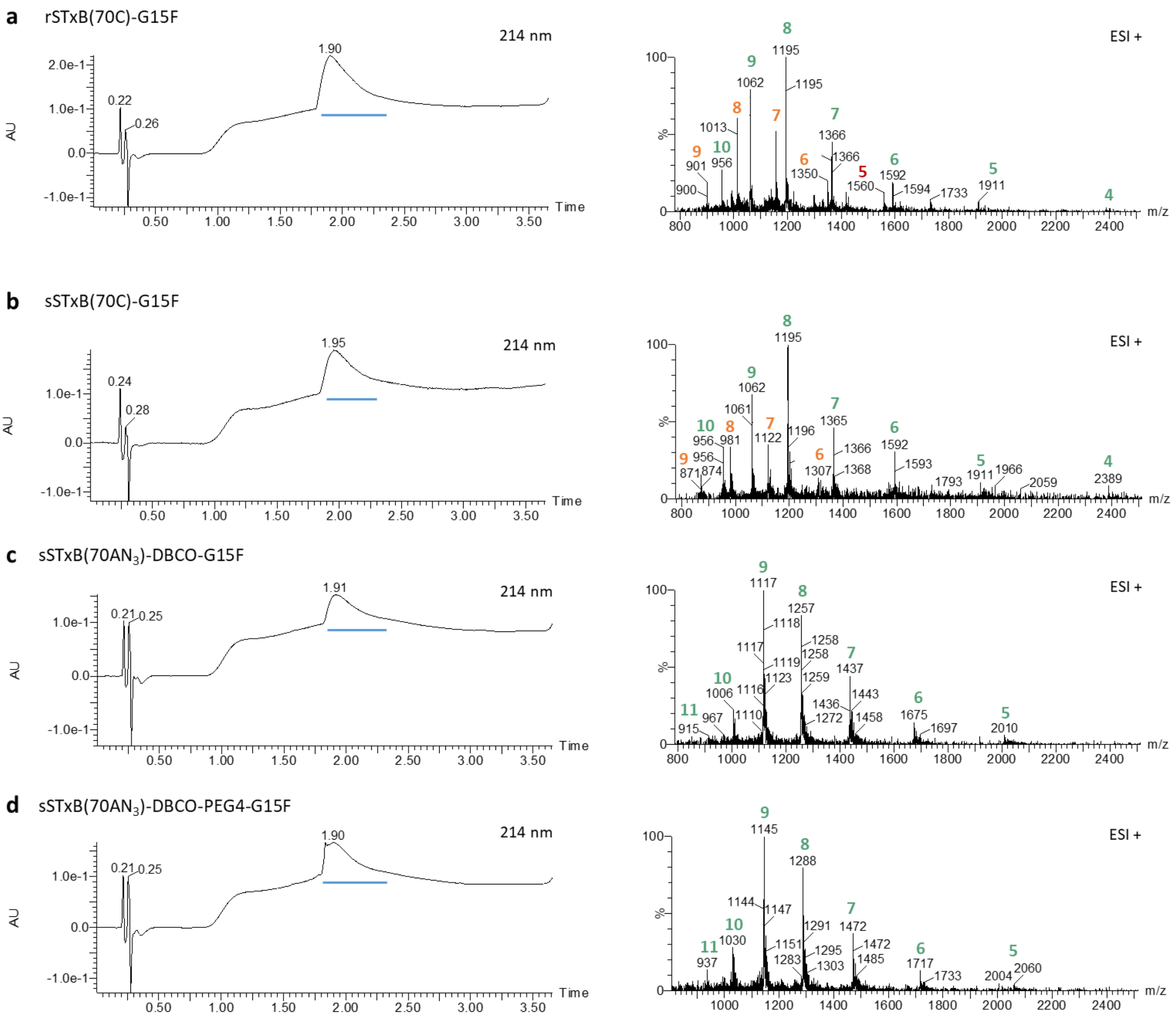

**Figure S18:** UPLC-MS analyses of sSTxB(70C) and sSTxB(70AN$_3$) conjugates with G15F.

Mass spectra (ESI+): Peaks corresponding to the expected product are annotated with their corresponding positive charge in green. Peaks annotated in orange correspond to monomers with a non-reactive cysteine (either due to a glutathione adduct on recombinant STxB, or to a *t*Bu adduct on synthetic STxB). Peaks annotated in red correspond to unconjugated reactive monomers.

a) $C_{420}H_{646}N_{114}O_{133}S_4$ $[M+4H^+]^{4+}$ calculated: 2389.4, found: 2387.6; $[M+5H^+]^{5+}$ calculated: 1910.9, found: 1911.3; $[M+6H^+]^{6+}$ calculated: 1592.6, found: 1592.4; $[M+7H^+]^{7+}$ calculated: 1365.2, found: 1365.5; $[M+8H^+]^{8+}$ calculated: 1194.7, found: 1194.6; $[M+9H^+]^{9+}$ calculated: 1062.1, found: 1062.1; $[M+10H^+]^{10+}$ calculated: 956.0, found: 956.2.

b) $C_{420}H_{646}N_{114}O_{133}S_4$ $[M+4H^+]^{4+}$ calculated: 2389.4, found: 2388.2; $[M+5H^+]^{5+}$ calculated: 1910.9, found: 1910.8; $[M+6H^+]^{6+}$ calculated: 1592.6, found: 1592.7; $[M+7H^+]^{7+}$ calculated: 1365.2, found: 1365.4; $[M+8H^+]^{8+}$ calculated: 1194.7, found: 1194.5; $[M+9H^+]^{9+}$ calculated: 1062.1, found: 1061.8; $[M+10H^+]^{10+}$ calculated: 956.0, found: 956.7.

c) $C_{446}H_{671}N_{120}O_{138}S_4$ $[M+5H^+]^{5+}$ calculated: 2010.6, found: 2009.9; $[M+6H^+]^{6+}$ calculated: 1675.7, found: 1675.2; $[M+7H^+]^{7+}$ calculated: 1436.4, found: 1436.6; $[M+8H^+]^{8+}$ calculated: 1257.0, found: 1256.9; $[M+9H^+]^{9+}$ calculated: 1117.5, found: 1117.5; $[M+10H^+]^{10+}$ calculated: 1005.8, found: 1005.8; $[M+11H^+]^{11+}$ calculated: 914.5, found: 915.0.

d) $C_{457}H_{692}N_{121}O_{143}S_4$ $[M+5H^+]^{5+}$ calculated: 2060.1, found: 2060.5; $[M+6H^+]^{6+}$ calculated: 1717.0, found: 1716.8; $[M+7H^+]^{7+}$ calculated: 1471.8, found: 1471.7; $[M+8H^+]^{8+}$ calculated: 1287.9, found: 1288.5; $[M+9H^+]^{9+}$ calculated: 1144.9, found: 1144.9; $[M+10H^+]^{10+}$ calculated: 1030.5, found: 1030.4; $[M+11H^+]^{11+}$ calculated: 936.9, found: 937.3.

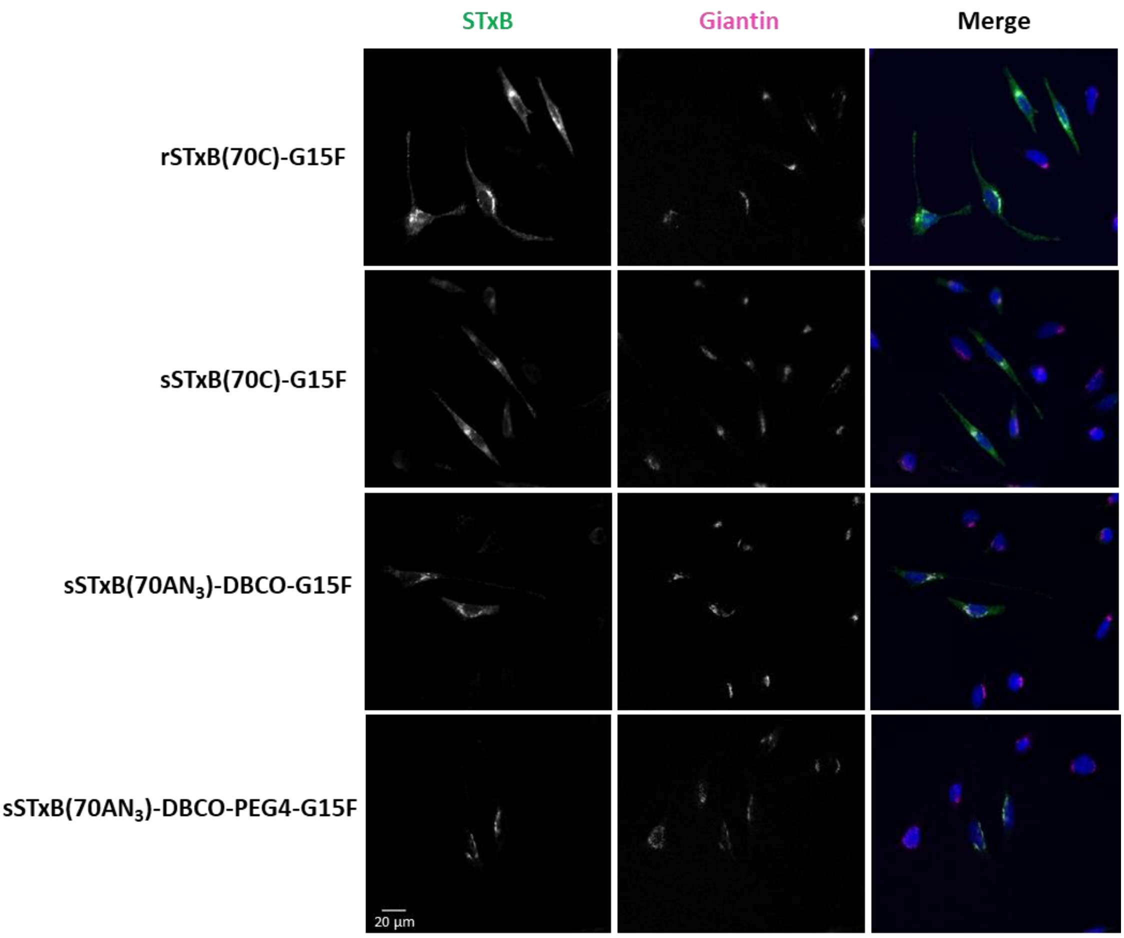

**Figure S19:** Retrograde trafficking on HeLa cells of sSTxB(70C) and sSTxB(70AN$_3$) conjugates with G15F. Cells were incubated with the indicated STxB conjugates for 30 min at 4 °C, followed by PBS washes and incubation for 50 min at 37 °C for synchronized internalization. Merge: STxB in green, Golgi in magenta, and DNA dye Hoechst in blue. Scale bar: 20 µm.
